# Bacteria break through one-micrometer-square passages by flagellar wrapping

**DOI:** 10.1101/2025.02.12.637796

**Authors:** Aoba Yoshioka, Yoshiki Y. Shimada, Toshihiro Omori, Naoki A. Uemura, Kazutaka Takeshita, Kota Ishigami, Hiroyuki Morimura, Maiko Furubayashi, Tetsuo Kan, Hirofumi Wada, Yoshitomo Kikuchi, Daisuke Nakane

## Abstract

Confined spaces are omnipresent in the micro-environments, including soil aggregates and intestinal crypts, yet little is known about how bacteria behave under such conditions where movement is challenging due to limited diffusion. Stinkbug symbiont *Caballeronia insecticola* navigates a narrow gut passage about one micrometer in diameter to reach the stinkbug’s symbiotic organ. Here, we developed a microfluidic device mimicking the host’s sorting organ, wherein bacterial cells are confined in a quasi-one-dimensional fashion, and revealed that this bacterium wraps flagellar filaments around its cell body like a screw thread to control fluid flow and generate propulsion for smooth and directional movement in narrow passages. Physical simulations and genetic experiments revealed that hook flexibility is essential for this wrapping; increasing hook rigidity impaired both wrapping motility and infectivity. Thus, flagellar wrapping likely represents an evolutionary innovation, enabling bacteria to break through confined environments using their motility machinery.

## Main Text

Bacteria live everywhere on earth, from deep-ocean trenches to the tops of mountains, and even inside animals and plants. The micro-habitats that bacteria inhabit are replete with confined environments such as small pores, cavities, and narrow channels^1^. Soil, for example, is a mixture of particles of various sizes at the micron scale, which creates complex confined spaces^2^. Interspaces between cells and microvilli are also confined^3^, where bacteria inhabit or invade. In recent years, bacterial behavior in micro-environments has been studied in some model species by using microdevices that mimic their natural habitats^4^. However, it remains largely unknown how environmental bacteria with diverse styles of motility behave in confined environments.

Bacterial motility adapts to fluid dynamics at low Reynolds number, where inertia is negligible and viscous forces are dominant^5^. Moreover, in confined spaces, the apparent viscosity increases dramatically at the solid-liquid interface due to interactions between materials at a boundary. From this physical viewpoint, moving in micrometer confined spaces, which bacteria often encounter in their natural habitats, is thought to be challenging for bacteria, like rock excavation.

A few notable examples of confined-space niches encountered by bacteria are found in animals, such as squids and insects, which sort specific bacterial symbionts from environmental microbiota by using a constricted region (CR) in front of their symbiotic organs. Such a sorting gate has been found in the squid-*Vibrio* luminescent symbiosis^6^, and recently well-described in the bean bug *Riptortus pedestris* which harbors with *Caballeronia insecticola* in its gut^7,8^. The structure of these sorting organs is very simple, a tubular structure with a lumen of one micrometer in diameter and a few hundred micrometers in length^9,10^, but is remarkably effective at sorting out the hosts’ specific symbionts only. In both cases the symbionts described above possess polar flagella, which are crucial for crossing their host’s CR, a narrow passage filled with a mucus-like matrix^10,11^. While bacteria commonly swim by rotating helical flagellar filaments, which trail the cell body^12^, recent studies have demonstrated a novel mode of swimming in these symbiotic bacteria wherein symbionts wrap flagellar filaments around the cell body and propel themselves with the rotary motor positioned at the front of the cell^13,14^. Since flagellar wrapping is known to be effective in moving in viscous conditions and escaping substrate surfaces^15^, this unique swimming mode is thought to play a pivotal role in adapting to the internal environments of the hosts^3^. Little is known, however, about the ecological significance of flagellar wrapping in spatially-confined narrow passages.

### Flagellar wrapping in the host’s sorting organ

We first investigated how the insect symbiont, *C*. *insecticola*, passes through the sorting organ of its host, *R. pedestris* (Fig. 1a,b). A second instar nymph was dissected after being fed *C. insecticola* expressing a green fluorescent protein (GFP), and the digestive tract was observed under optical microscopy around the entry gate of the symbiotic organ, CR (Fig. 1c,d). *C. insecticola* cells located by GFP signal had crossed the CR and were observed arranged in a row at the subsequent narrow passage connecting the M4B and M4 midgut regions 2-4 hours after feeding. Time-lapse imaging revealed that the cells showed repeated back-and-forth movements, but moved directionally from the M3 to M4 regions through the CR with a net displacement of 50 μm min^−1^ (Fig. 1e and Movie S1), indicating that *C*. *insecticola* cells can break through the host’s narrow passages, a 200 μm distance, in a few minutes. Flagellar filaments of *C*. *insecticola* can be fluorescently labeled with amine-reactive dyes^13^. High-speed imaging of *C*. *insecticola* with fluorescently-labelled flagellar filaments revealed that *C*. *insecticola* adopts a wrapped mode of flagellar motility while in the CR-M4B narrow passage (Fig. 1f,g and Movie S2), suggesting that flagellar wrapping is important for breaking through the host’s sorting organ.

**Fig. 1.**
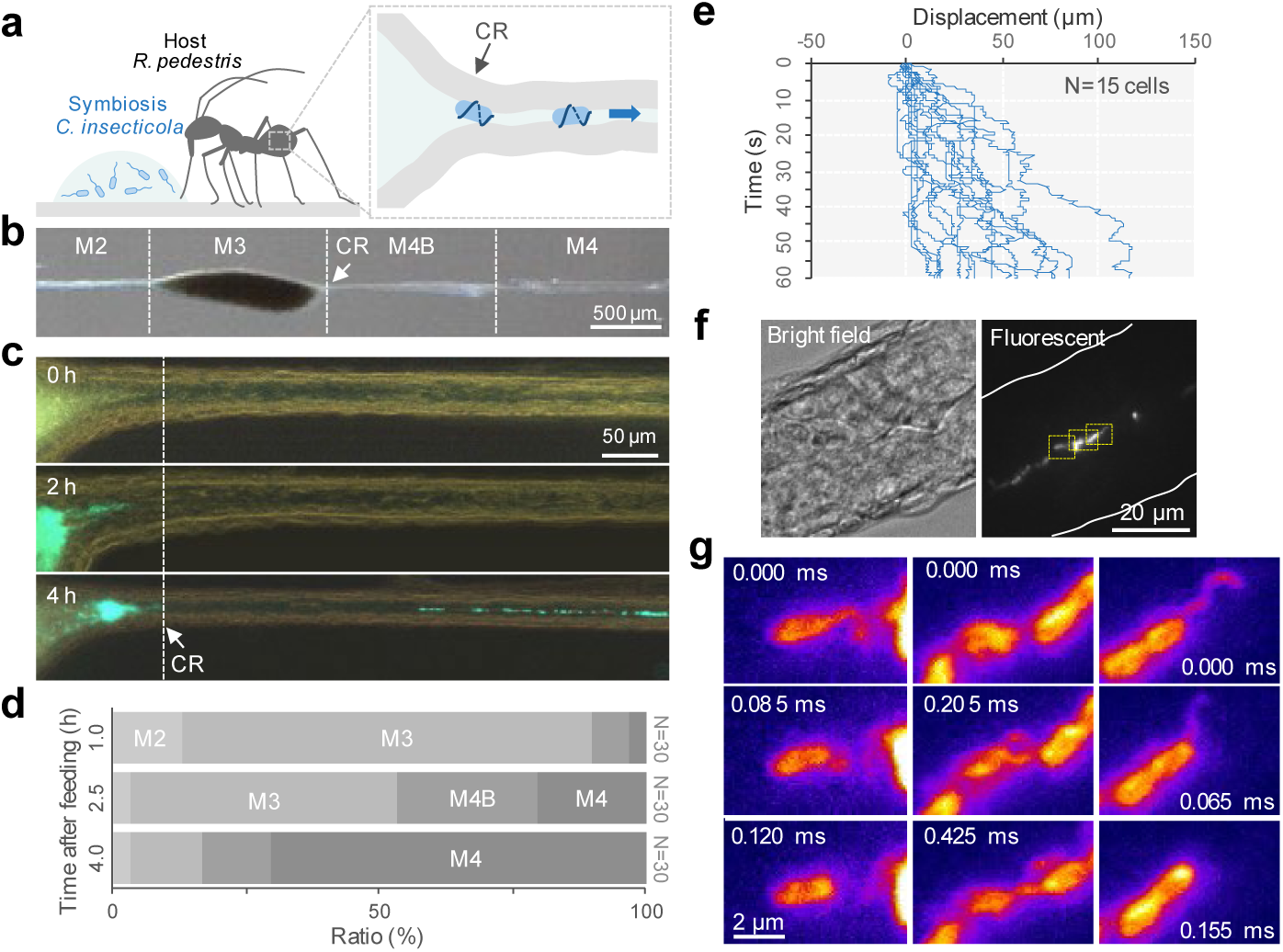
Single cell imaging of flagellar wrapping motility in the gut sorting organ. **a**, Schematic illustration of a second instar nymph of *R. pedestris* fed with *C. insecticola*. **b**, Midgut organization and the bacterial sorting organ (CR). **c**, Symbiont localization at the CR-M4B region. The images show the gut region 0, 2, and 4 hours after feeding of GFP-labeled *C. insecticola*. Note the symbionts start entering the CR 2h after feeding and are passing through the narrow duct 4h after feeding. See also Movie S1. **d**, Time required for symbiont sorting. Ratio of the locations where the GFP signal from *C. insecticola* was detected at the most distal part is shown (N = 30 nymphs). **e**, Time course of symbiont displacement in the CR-M4B region 2-4 hours after feeding of GFP-labeled *C. insecticola*. Single-cell movement colored by a blue line, and 15-cell displacements are overlayed. **f**, Direct visualization of the flagellar filament of the symbiont in the M4B region. *C. insecticola* cells fluorescently labeled by amine-reactive dye were fed to a second instar nymph, and the gastrointestinal tract was dissected 2-4 hours after infection. *Left*: Bright-field. *Right*: Fluorescent. See Movie S2. **g**, Flagellar wrapping in the M4B region. Yellow-boxed regions in **f** were magnified, and time-lapsed images are presented.

### Our device: quasi-one-dimensional device (Q-1D) mimicking the sorting organ

To observe the symbiont’s behavior in micrometer-level narrow passages in more detail, we created a dimethylpolysiloxane (PDMS) microfluidic device with linear, open channels whose width and depth were limited to 1 µm, and filled with a viscous solution of 0.4% methylcellulose (MC) (Fig. S1a,b), which accurately mimics the physical parameters of the insect’s sorting organ. In this device, the diffusion of microbeads was almost entirely restricted, with their apparent diffusion coefficient being reduced to 1/60 of that in a standard sample chamber (Fig. S1c-e). When *C. insecticola* cells were confined in the device with MC (Fig. 2a), the cells oriented themselves parallelly along the narrow passages and exhibited directional movement (Fig. 2b,c and Movie S3), whereas they moved in a directionally unbiased manner within the standard chamber. In this device, the symbiont cells showed frequent switchbacks within a short time period of a few seconds but were biased to one side for a longer period of 1 minute at a net displacement of 80 μm min^−1^ (Fig. 2d). The time from one directional change to the next showed a distribution of single exponential decay in both forward and backward directions (Fig. 2e), whereas the bias in the forward direction gradually increased with time (Fig. 2f). Considering that the deletion mutant of *cheA* stopped movement in the device for a long time (Fig. S2 and Movie S4), clockwise (CW) rotation of the flagellar motor is involved in directional movement. We used this quasi-one-dimensional device (Q-1D) for the remainder of this study.

**Fig. 2.**
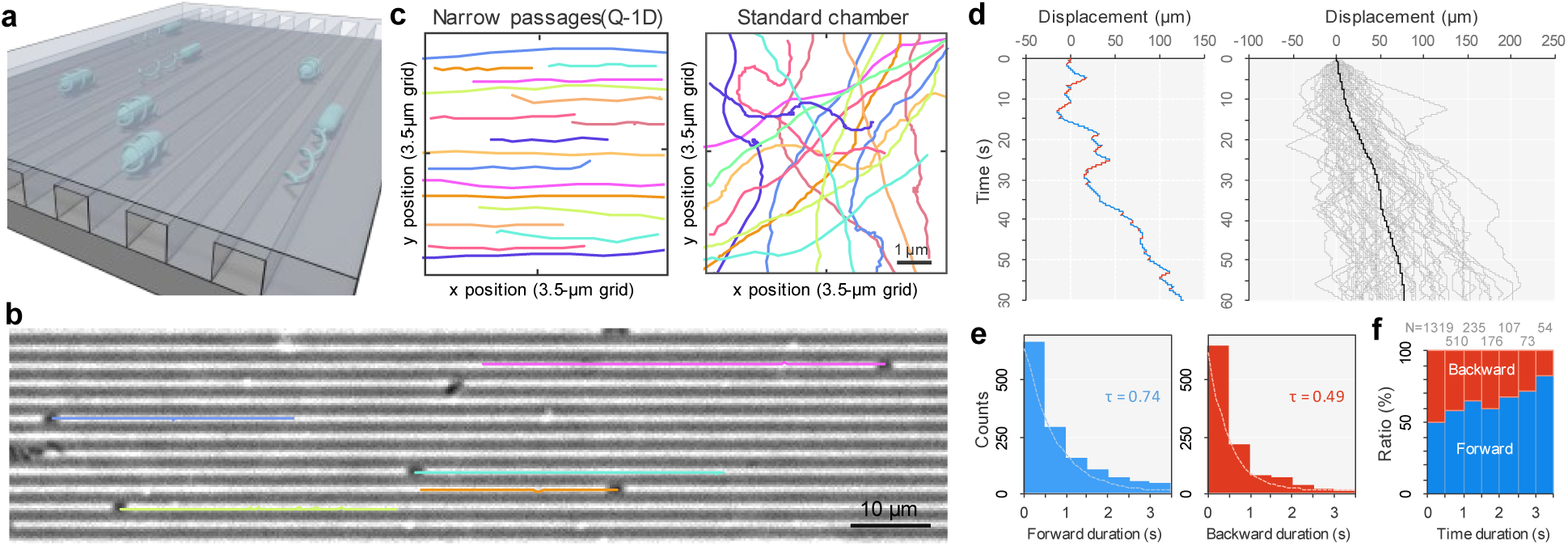
Smooth and directional movement of symbiotic bacteria in microfluidic narrow passages. **a**, Schematic of confined bacteria in the Q-1D microfluidic device. **b**, Smooth and directional movement of *C. insecticola* cells in Q-1D. The cell trajectories over 20 s were overlayed on the phase-contrast image (see Movie S3). **c**, Single-cell trajectories for 1 s. *Left*: Q-1D. *Right*: Standard chamber. **d**, Time course of cell displacement in Q-1D. *Left*: Single cell. Forward and backward movements were detected as positive and negative displacements along the x axis, and presented by the blue and red colored lines, respectively. *Right*: Overlays of 50 cells and the average. **e**, Distribution of the forward and backward duration of swimming time in Q-1D. Duration from one directional change to the next were measured and color-coded as in **d**. Dashed lines show the fit of single exponential decay, where time constant τ is presented. **f**, Duration ration of forward and backward direction. The ratio was sorted by 0.5 s, and the ratio at each time point is presented.

### Flagellar wrapping gives an advantage in narrow passages

The Q-1D also allowed us to observe the motility of *Salmonella enterica*, a model species for the study of bacterial flagella (Fig. S3a, Fig. S4a and Movie S5). However, the directionality of their movement in the narrow passage was less clear (Fig. S3b). The time intervals between directional changes in *S. enterica* followed a single exponential decay, with a time constant comparable to that observed in *C. insecticola* (Fig. S3c), whereas the forward directional bias remained relatively consistent over time (Fig. S3d). In contrast, *Vibrio fischeri* (*Aliivibrio fischeri*), the luminescent symbiont of the Hawaiian bobtail squid *Euprymna scolopes* ^6^, moved smoothly in Q-1D (Fig. S4a and Movie S6). Remarkably, the average of mean square displacement (MSD) of *V. fischeri* and *C. insecticola* in Q-1D showed parabolic curve, as well as that of *C. insecticola* in CR-M4B narrow passage in host, indicating directional movement, while that of *S. enterica* in Q-1D increased linearly, suggesting random movement (Fig. S4b). This difference may be caused by the arrangement of the motility machinery, *i.e.* multiple flagella randomly distributed over the entire cells in *Salmonella*, while a single or few flagella extend from one end of the cell in *C*. *insecticola*, and *V*. *fischeri*, respectively^15^, or due to the lack of flagellar wrapping in *Salmonella*. To further investigate the importance of polar flagella in navigating confined spaces, we observed motility in bacterial species closely related to *C. insecticola* and possessing polar flagella using the Q-1D device.

In total, 13 species of *Burkholderia* sensu lato and allied groups, including members of the genera *Caballeronia*, *Paraburkholderia*, *Burkholderia*, *Pandoraea*, and *Cupriavidus*, were compared (Fig. 3a). While all these species swam freely at 15-34 μm s^−1^ on average in the standard chamber (Fig. 3b), their movements in Q-1D showed a clear difference (Fig. 3c). Eight species, including *C*. *insecticola* and *Pandoraea norimbergensis*, exhibited smooth and directional movement with a net displacement of 40-130 μm min^−1^ on average and 3-10% of their average speed in the standard chamber (Table S1 and Movie S7). In contrast, five species including *B. anthina* were barely able to move in Q-1D with a displacement of 0-6 μm min^−1^, a decrease to 0-0.4% of their instantaneous speed in the standard chamber (Movie S8). These results demonstrated that ability to move through confined passages varies even in closely related bacterial species carrying polar flagella. Notably, ability to move within confined spaces positively correlated with infection rates of these bacterial species in the bean bug host (Fig. 3d), with bacteria showing smooth and directed movement in Q-1D having an almost 100% infection rate while bacteria with poor movement in Q-1D showing no infection.

**Fig. 3.**
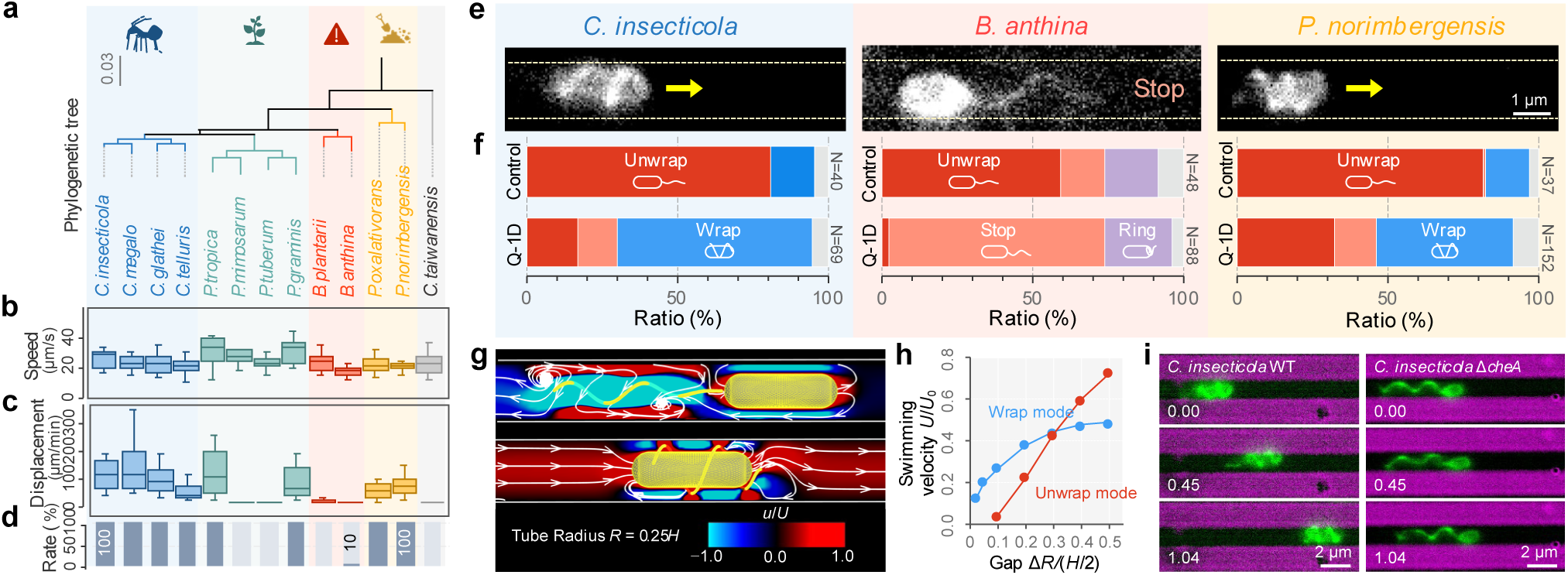
Interspecies variation in flagellar wrapping and its advantages in confined spaces. **a**, Phylogenetic tree of *Burkholderia sensu lato* group and *S*. *enterica* based on 16S ribosomal RNA sequences. Stinkbug-associated *Caballeronia*, plant-associated *Paraburkholderia*, pathogenic *Burkholderia* sensu stricto group, and *Pandoraea* species are highlighted with blue, green, red, and yellow, respectively^7^. **b**, Swimming speed for 1.0 s in growth medium for each species. **c**, Cell displacement for 1 min in Q-1D. Box plots in **b-c** represent the minimum, maximum, sample median, and the first and third quartiles (N = 20 cells). **d**, Infection ratio of the bean bug *R. pedestris* (data derived from our previous paper^7^). **e**, Fluorescently labeled flagellar filament of *C. insecticola*, *B. anthina*, and *P. norimbergensis* in Q-1D. The edge of the narrow passage is presented by dashed yellow lines. The arrows indicate the moving direction of each cell. See Movie S10. **f**, The fraction of flagellar filament morphology in growth medium and Q-1D. The wrapped, unwrapped, incompletely wrapped, and uncategorized flagella are colored by red, purple, blue, and gray, respectively. **g**, Numerical calculations of time-averaged flow field around the bacterial model in a narrow tube. *Left*: Unwrapped mode. *Right*: Wrapped mode. White arrows are the streamline and contour color indicate the velocity component in the swimming direction normalized by the swimming velocity *U*. **h**, Swimming velocity *U* as a function of the gap in the cylinder tube Δ*R*. The values are normalized by the swimming velocity of the unwrapped mode in the free space *U*_0_ and a half body length *H*/2, respectively. **i**, Snapshot of cell behavior in Q-1D. Fluorescently labeled flagellar filament of *C. insecticola* WT and Δ*cheA*. See Movie S11.

To clarify why there is such a difference in movement between closely related species with polar flagella, we visualized how they use their flagellar filaments within standard chamber and Q-1D by amine-reactive dye staining^13^. Among the 13 species, we were able to visualize the flagellar filaments of four species, *C*. *megalochromosomata*, *B*. *anthina*, *P*. *norimbergensis* and *P*. *oxalativorans,* as in *C. insecticola* (Table S1). High-speed imaging revealed that *C*. *megalochromosomata*, *P*. *norimbergensis* and *P*. *oxalativorans* wrapped their flagellar filaments around their cell body during switchbacks under high viscosity (Fig. S5a and Movie S9), as seen in *C*. *insecticola*^13^. The helical shapes of the flagellar filaments were consistent with normal and coil forms in polymorphic transformations^16^, changing from 2 to 1 μm in helix pitch and from 0.3 to 0.5 μm in helix radius after flagellar wrapping (Fig. S5b). However, *B*. *anthina* exhibited no flagellar wrapping (Fig. S5a and Movie S9); the helical filaments folded longitudinally proximal to the cell surface in an apparent ring with larger diameter than that found in the normal form. Furthermore, flagellar wrapping was observed at a high frequency in *C*. *insecticola* and *P*. *norimbergensis* in Q-1D, while no wrapping was observed in *B*. *anthina*, as in the standard chamber, (Fig. 3e and Movie S10). Notably, in *C*. *insecticola* and *P*. *norimbergensis* the ratio of wrapped cells in Q-1D increased to 65%, 3-4 times higher than in the standard chamber (Fig. 3f). These results suggest that these bacterial species, presumably triggered by spatial limitations, change their swimming style to be biased toward flagellar wrapping to achieve efficient locomotion in narrow passages, resulting in directional movement.

We used numerical simulations to interrogate why flagellar wrapping is dominant in narrow passages. A fluid-mechanical model of a swimming bacterium shows two distinct models of unwrapped and wrapped flagella mode (Fig. S6a), and the bacterial models were considered to be swimming unidirectionally in a narrow tube (Fig. S6b and see more details in the SI text). In contrast to the flow field in free space (Fig. S6c), no lateral suction flow is generated by the unwrapped flagellum in the narrow tube (Fig. 3g *Top*). The fluid friction between the cell body and the wall provides strong resistance such that the flow only contributes to the agitation of the fluid around the flagellum. On the other hand, the wrapped flagellum scrapes the fluid in the gap like a corkscrew, creating a laminar flow structure in the narrow tube and contributing to cell propulsion (Fig. 3g *Bottom*, and Fig. S6c). This seems to explain our observations of cells swimming within Q-1D (Fig. 3i and Movie S11). The speed of the unwrapped mode decreases monotonically as the gap between the wall and the bacterium becomes narrower (Fig. 3h *Red*). However, the swimming speed of wrapped-mode cells is maintained even in the narrow tube (Fig. 3h *Blue*). Assuming a cell length of *H* = 2.5 μm, the gap in a circular tube of 1 μm diameter is Δ*R*/(*H*/2) = 0.025, and the swimming speed is estimated to be *U* = 180 μm/min for the wrapped mode, which is comparable to the observed speed of *C. insecticola* (Table S1). The effect of the gap on the rotational angular velocity is relatively small in both modes (Fig. S6d), indicating that the flagellar motor produces sufficient torque to generate cell rotation even in a narrow channel. These results confirm that there is a physical advantage of flagellar wrapping in narrow passages, and therefore wrapped-mode swimming is likely dominant in such confined environments.

### Moderately flexible hook for flagellar wrapping

We wondered which flagellar component/feature is responsible for wrapping motility. In *Campylobacter* and *Shewanella*, which exhibit wrapping motility, flagellar filaments are composed of two types of flagellin monomers (FlaA/FlaB), and it is thought that the flagellin comprising the cell-proximal portion of the flagellar filament provides the flexibility for the flagellar filament to wrap around the cell body^17,18^. Since *C*. *insecticola* has only one flagellin gene (*fliC*)^19^, we focused on the hook, the 60 nm long structure that connects the flagellar motor to the filaments^12,20^, as a candidate that could contribute to the flexibility of the flagellar filament required for wrapping. When cells were immobilized on a glass slide and flagellar rotation was inactivated by carbonyl cyanide 3-chlorophenylhydrazone (CCCP), the filaments at the cell-proximal end wobbled at a small angle due to Brownian motion (Movie S12). Based on this observation, the thermal fluctuation of fluorescently labeled flagellar filaments was measured as the flagellar orientation angle at 5-ms time resolution (Fig. S7), the variances of the orientation angle (*σ* = 0.038 ± 0.008 for *C. insecticola* and *σ* = 0.018 ± 0.006 for *B. anthina*), were used to obtain the hook bending stiffness, *A_hook_*_12_. Assuming that the number of flagellar filaments is three, the stiffness of *C*. *insecticola* (*A_hook_* = 0.67×10^−2^^5^ Nm^2^) is 5 times lower than that of *B. anthina* (*A_hook_* = 0.15×10^−2^^5^ Nm^2^) (Supplementary Text), consistent with the previously reported value measured under a low torque regime in the flagellar motor of *Escherichia coli*^20^.

Next, we constructed a mechanical model of flagellar wrapping, in which the flagella bundle is regarded as a single elastic helical filament with short-range attractive interactions (Fig. S8a and SI text). For the deformations of a flexible filament, we computed all the elastic modes of stretching, bending, and twisting, taking full account of the geometric nonlinearities inherent in such slender structures, based on a previously reported method^21^. Three qualitatively distinct wrapping behaviors were observed in simulations with varying hook/filament stiffness ratios, *A*_hook_/*A*, and filament twist/bend stiffness ratios, *C*/*A* (Table S2 and Fig. S8b). A flagellar filament with a flexible hook at *A*_hook_/*A* = 0.02 and *C*/*A* = 0.75 showed a smooth transition to wrapping around the cell body (Movie S13), as observed in *C. insecticola* (Fig. 4a *Left*). On the other hand, a filament with a more rigid hook at *A*_hook_/*A* = 0.14 and *C*/*A* = 0.75 performed incomplete wrapping that overlapped near the end of the cell body (Movie S14). For an even smaller twist/bend stiffness ratio at *C*/*A* = 0.5, this feature was more evident, where the filament folded into a ring at the cell-proximal end without wrapping at all (Movie S15), as we observed in *B. anthina* (Fig. 4a *Right*, and see also Movie S16). Our mechanical modeling and numerical simulations suggest that flagellar wrapping can be explained by a single factor, the stiffness of the hook in polar flagellated bacteria.

**Fig. 4.**
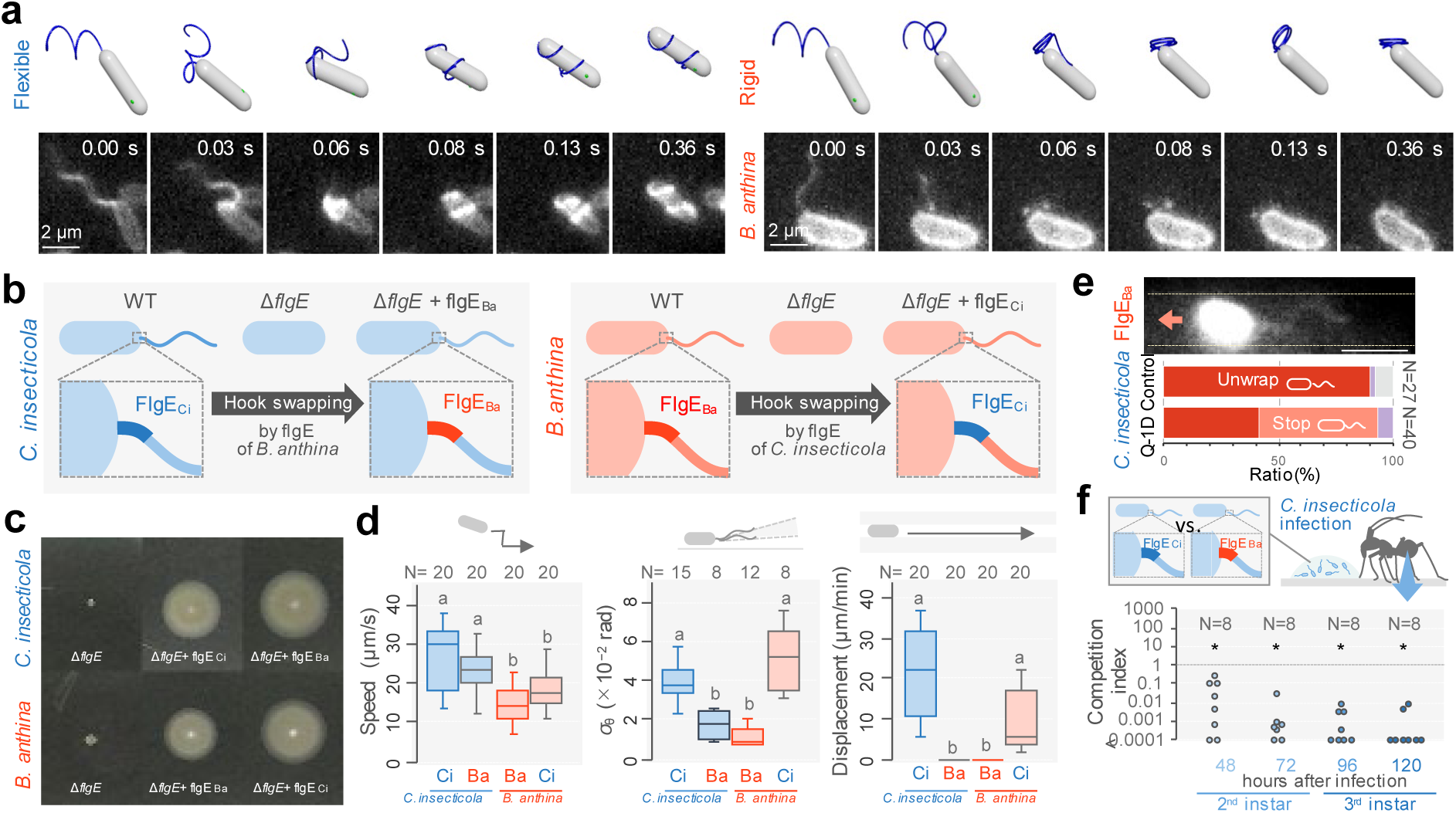
Moderately flexible hook for flagellar wrapping. **a**, Sequential images of the flagellar filament during cell switchbacks. *Upper*: Numerical calculation of the flexible and rigid hook. The coil form of flagellar filaments is transformed by the CW rotation. *Lower*: Fluorescent images of *C. insecticola* and *B. anthina*. See also Movie S16. **b**, Schematic of the genetic swapping of hook by *flgE* deletion and transformation. **c**, Motility assay on soft agar of the *flgE* swapping mutants. **d**, Characterization of *flgE* swapping mutants. *Left*: Swimming speed for 1.0 s in growth medium. *Center*: Variance of the flagellar orientation angle *σ*. *Right*: Cell displacement for 1 min in Q-1D. Box plots present the minimum, maximum, sample median, and the first and third quartiles. Schematic of the measurements is presented at the top of each graph. Different letters indicate statistically significance (P < 0.05). Statistical significance was analyzed by the Mann– Whitney U test with Bonferroni correction. **e**, Direct visualization of flagellar filaments in the *flgE* swapping mutants. *Top*: Fluorescent image in Q-1D. The edge of the narrow passages is represented by dashed lines. The arrows indicate the direction of movement of the cell. *Bottom*: Fraction of flagellar filament morphology in growth medium and Q-1D. The color codes of bars are the same as in Fig. 3f. **f**, *In vivo* competition assay. Infection competitiveness of *C. insecticola* with FlgE_Ci_ against *C. insecticola* with FlgE_Ba_ in the gut symbiotic organ of *R. pedestris*. The competitive index (CI) was calculated by (output *C. insecticola* with FlgE_Ba_ colony count/input *C. insecticola* with FlgE_Ba_ colony count)/(output C*. insecticola* with FlgE_Ci_ colony count/input *C. insecticola* with FlgE_Ci_ colony count). One sample Wilcoxon tests were conducted for each CI (against CI = 1), (*P* < 0.05).

To test this model, we constructed hook-swapping mutants by expressing the hook component protein FlgE of *B. anthina* (FlgE_Ba_) in *C. insecticola* and FlgE of *C. insecticola* (FlgE_Ci_) in *B. anthina* (Fig. 4b,c). The swimming speed of both swapping mutants was almost the same as that of their wildtype (Fig. 4d *Left*), while the *σ* was measured to be 0.011 for *C. insecticola* with FlgE_Ba_ and 0.051 for *B. anthina* with FlgE_Ci_ (Fig. 4d *Center*). We found *C. insecticola* with FlgE_Ba_ decreased cell displacement in Q-1D to almost zero, whereas *B. anthina* with FlgE_Ci_ increased its cell displacement in Q-1D to the same level as wildtype *C. insecticola* (Fig. 4d *Right*). Indeed, *B. anthina* with FlgE_Ci_ started to perform flagellar wrapping (Movie S17), although infrequently, potentially due to the swapped hook being too flexible to fully replicate the behavior of wildtype *C. insecticola*. In contrast, *C. insecticola* with FlgE_Ba_ was no longer able to wrap their flagellar filaments around their cell bodies in Q-1D (Fig. 4e) and showed a strong reduction of their infection competitiveness in the host *R. pedestris* (Fig. 4f). Together, these results demonstrate a pivotal role of the hook not only in the flagellar wrapping but also in cell displacement in Q-1D and in insect-microbe symbiosis.

## Discussion

We investigated the physical and ecological role of flagellar wrapping in insect symbionts, revealing that this mode of flagellar motility gives bacteria an advantage in confined spaces (Fig. 2 and Fig. 3) and promotes the establishment of symbiosis (Fig. 1 and Fig. 4). In addition to insect symbionts and other symbionts such as *V. fischeri*^13,22^, serious human pathogens such as *Campylobacter jejuni*, *Helicobacter* spp., and *Pseudomonas aeruginosa* are also known to exhibit flagellar wrapping^18,23,24^. Therefore, it is probable that flagellar wrapping is a common strategy for infecting host animals when viscous conditions, such as mucous and confined regions like gut lumen crypts, are encountered. Moreover, this motility could also be beneficial in complex constricted spaces such as soil, opening a new window to understand the ecology of environmental bacteria.

Although we were not able to visualize the flagella of all tested bacteria, considering that the directional motility in Q-1D is due to flagellar wrapping (Fig. 2), this characteristic swimming style may be common in *Burkholderia* sensu lato. The flagellar wrapping in *Pandoraera*, the sister group of *Burkholderia* sensu lato, suggests that this motility was developed by the common ancestor of these bacterial groups, and *B. anthina* reverted to an unwrapped swimming style (Fig. 2 and Fig. S5). From an evolutionary perspective, it would be of great interest to understand the selective forces that have led some species to wrap while other closely related species do not. This study demonstrated that while hook flexibility is important for flagellar wrapping; however, this flexibility may cause inefficient rotation in the normal unwrapped mode of flagellar motility. If so, there may be a trade-off between the wrapped and unwrapped motility and, in some environmental niches, flagellar wrapping and hook flexibility may be disadvantageous.

Another notable point is that wrapping motility could be induced by a mechanical stimulus in narrow passages (Fig. 3 and Fig. 4). In *V*. *fisheri*, cells alter their behavior upon entry into a confined space, allowing them to escape from confinement^25^. In *Shewanella putrefaciens*, flagellar wrapping enables efficient movement away from a substratum using instantaneous CW rotation of the flagellar motor^14,15^. Similarly, in *E. coli*, CW bias gradually increases under high load^26^. Wrapped-mode bias in narrow passages may be explained by a universal physical process such as the loads triggered by spatial limitations and intrinsic properties of the flagellum itself. Alternatively, wrapped-mode bias could be due to an unknown physiological pathway in bacteria that senses ambient physical conditions.

Since the rotational model of the bacterial flagella was established in 1974^27–29^, the machinery and mechanisms of flagella have been widely investigated, but flagellar wrapping is a much more recent discovery. Although the number of culturable bacteria is still extremely limited, a wide variety of movement patterns and machinery have been found in bacteria^5^, but their ecological significance and evolutionary process are still largely unknown^30^. Mimicking their micro-habitat in microfluidic devices in conjunction with analyzing the physical properties and genetic backgrounds of different species, as shown in this study, will reveal new bacterial behaviors and adaptive strategies in natural environments.

## Acknowledgments

We thank Eli Cohen, Daisuke Takagi, Jonathan Lynch, and Peter Mergaert for critical reading and comments on the manuscript.

## Funding

This study was supported partly by KAKENHI grants from the Japan Society for the Promotion of Science (JSPS), and funds from Noguchi Institute and Precise Measurement Technology Promotion. The funders had no role in study design, data collection and analysis, decision to publish, or preparation of the manuscript.

JSPS KAKENHI grant 22H05066 (DN, TK)

JSPS KAKENHI grant 22H05067 (HW)

JSPS KAKENHI grant 22H05068 (YK, KT)

Foundation from Noguchi Institute (DN)

Foundation from Precise Measurement Technology Promotion (DN)

## Author contributions

Conceptualization: HW, TK, YK, DN, Methodology: All authors, Formal analysis and investigation: All authors, Funding acquisition: HW, YK, DN, Writing – original draft: AY, WH, YK, DN, Writing – review & editing: YK, DN

## Competing interests

Authors declare that they have no competing interests.

## Data and materials availability

All data are available in the main text or the supplementary materials.

## Supplementary Materials

Materials and Methods

Supplementary Text

Figs. S1 to S8

Tables S1 to S5

References

Movies S1 to S17

## Supplementary Materials

### Materials and Methods

#### Strain and culture conditions

*C. insecticola* (former called *Burkholderia insecticola*) RPE64 cells and other bacteria belongs to *Burkholderia sensu lato* group were grown to an early log phase in yeast-glucose (YG) liquid medium or agar plate [0.5% (wt/vol) yeast extract, 0.4% (wt/vol) glucose and 0.1% (wt/vol) NaCl] at 28 °C^7^. *S. enterica* was grown in LB liquid medium [1% (wt/vol) tryptone, 0.5% (wt/vol) yeast extract and 0.1% (wt/vol) NaCl] at 37 °C^31^. *V. fisheri* was grown to an early log phase in seawater tryptone (SWT) liquid medium [0.5% (wt/vol) tryptone, 0.3% (wt/vol) yeast extract, 0.3% (wt/vol) glycerol, 70% (vol/vol) artificial seawater] at 28 °C ^32^. Bacterial species used in this study and their culture conditions are listed in Table S3.

#### Construction of deletion mutants and complemented strains

Flagella hook gene, *flgE* (BRPE64_ACDS27220 of *C*. *insecticola* or BAN20980_05336 of *B*. *anthina*), was deleted by the homologous recombination-based method with the suicide vector pK18mobsacB^33^. The successful deletion was confirmed by Sanger sequencing. Gene complementation was performed with the broad-host-range vector pBBR122. The fragment of *flgE* gene and its upstream region from *C*. *insecticola* or *B*. *anthina* was cloned into the Nco1 site of pBBR122 with NEBuilder HiFi DNA Assembly Master Mix (New England Biolabs). The constructed vectors were then introduced into the deletion mutants by electroporation. Primers used in this study were listed in Table S4. Strains and mutants used in this study were listed in Table S5.

#### Microfluidic device

Microfluidic devices (Q-1D) were fabricated using standard photolithography and soft lithography methods as described previously^34^. Briefly, polydimethylsiloxane (PDMS, Sylgard 184, Dow), a two-part silicone elastomer, was cast over a photolithography master and cured at room temperature for 48 h. The photolithography master was prepared by a photoresist (OFPR 800 23cp, Tokyo Ohka, Japan) pattern on a flat silicon wafer beforehand. The photoresist was spin-coated with a speed of 4000 rpm for 60 sec to have ∼1 µm thickness, and photolithographically patterned to have 1.5 µm wide stripes. The PDMS casting and curing on the master replicates the stripe patterns, making microfluidic channels with similar dimensions with the master pattern in a negative/positive inverted manner. A piece of PDMS was cut out using a scalpel and used as a microfluidic device. Cell suspensions with YG medium containing 0.4-0.5% methylcellulose (M0512, viscosity: 4,000 cP at 2%, Sigma-Aldrich) were dropped onto a glass slide and then covered with the microfluidic device casting from the top, which traps bacteria in the channels. The cell behaviour was observed within 20 min after the confinement. Fluorescent beads (Cat No. F8811; Thermo Fisher) were diluted 1/200 in water containing 2% bovine serum albumin and 0.4% methylcellurose.

#### Phylogenetic tree

Genome data of representative species of the genus *Burkholderia sensu lato* and related beta-proteobacteria were downloaded from the NCBI Assembly database. The multiple alignments of the 16S rRNA gene were constructed using Clustal W (https://www.genome.jp/tools-bin/clustalw) and displayed by iTOL (https://itol.embl.de/).

#### Measurements of swimming speed

The cell culture was centrifuged at 10,000 × g for 4 min, and the pellet was suspended in fresh liquid medium. The sample chamber was assembled using a coverslip with two pieces of double-sided tapes to create a thin channel for observing bacterial swimming motility. The cell suspension was poured into the chamber, and both ends of the chamber were sealed with nail polish to keep the sample from drying.

#### Preparation of fluorescently labeled cell

The method was according to the previous report^13^. Cell culture of 250 μL was centrifuged at 10,000 × g for 1 min at 25 °C, and resuspended in 1 mL of 0.1 M sodium phosphate buffer (pH 7.5). The suspension was mixed with 25 μg of DyLight 488 NHS ester (Thermo Fisher) or Cy3 NHS ester (Lumiprobe) and incubated at room temperature for 10 min. The suspension was washed twice in the sodium phosphate buffer, and resuspended in the motility buffer (20 mM potassium phosphate buffer pH 6.0 and 20 mM glucose).

#### Measurement of hook flexibility

The surface of the coverslip for sample observation was pre-treated with collodion to immobilize the cell on the surface. The fluorescently-labeled cells suspended in phosphate-buffered saline (PBS), made with 75 mM sodium phosphate (pH 7.4) and 68 mM NaCl in the presence of 50 μM CCCP at the final concentration were poured into the chamber, and both ends were sealed with nail polish to keep the sample from drying. The bacteria that tightly attached to the glass surface were used for observation in fluorescent microscopy.

#### Optical microscopy

Bacterial movements in Q-1D and the chamber were visualized under a phase-contrast microscope (IX73, Olympus) equipped with a 20× objective lens (UCPLFLN20X, NA 0.70, Olympus), a CMOS camera (DMK 33UX174, Imaging Source), and an optical table (HAX-0806, JVI). Projections of the images were captured as greyscale images with the camera under 0.1-s resolution and converted into a sequential TIF file without any compression. All data was analyzed by ImageJ 1.53k (rsb.info.nih.gov/ij/) and its plugins, TrackMate^35^, and particle tracker^36^.

For direct visualization of fluorescently labeled cells in both the Q-1D and the host midgut, the sample was examined under an inverted microscope equipped with a ×100 objective lens (UPLXAPO100×OPH, NA1.45, Olympus), a dichroic mirror (Di02-R488, or Di02-R532, Semrock), and an emission filter (FF01-540/80, Semrock, or FELH0550, Thorlabs). A laser beam (488-nm or 532-nm wavelength, OBIS488 or OBIS532, Coherent) was introduced into the inverted microscope through a lens for epi-fluorescence microscopy. Projections of the images were captured with the camera under a time resolution of 5-10 ms.

To observe the symbionts in real-time within the host midgut, the bacterial cells in the midgut were observed under an inverted microscope equipped with a ×40 objective (LUCPLFN40X, NA 0.60, Olympus), a filter-set (59022, Chroma), and a mercury lamp (U-HGLGPS, Olympus). Projections of the images were captured with the camera under 0.2-s resolution.

#### Numerical calculation

All the code and data to reproduce the numerical results reported in this manuscript are available from the authors upon reasonable request. Visualizations of simulated configurations are made using a free ray-tracing software POV-Ray3.7.0 (Unofficial version for Intel Mac OS). A movie file is produced from a series of configuration images by the command FFmpeg.

#### Motility assay on soft agar plate

Colony of *C. insecticola* and *B. anthina* were spotted on soft agar plate containing 0.4% agar and YG medium. For preparation and cultivation of the mutant strains, antibiotics were added to the medium at the following concentrations: kanamycin 30 and 100 µg ml^−1^ for *C. insecticola* and *B. anthina*, and chloramphenicol 30 µg ml^−1^ for *B. anthina*. Spreading of bacteria on the plate was captured after 24 h of incubation at 28 °C.

#### Insects rearing and symbiont infection

The bean bug *R. pedestris* was reared in the laboratory under a long-day regimen (16 h light, 8 h dark) at 25 °C in petri dishes (90 mm in diameter, 20 mm high)^10^. Soybean seeds and ion exchanged water were provided to the bugs as food and water, respectively. Newly molted second instar nymphs were fasted from water overnight, and then fed with suspension of symbiotic bacteria expressing mCherry or GFP^37^ or labeled cell body and flagellar filaments by DyLight 488 NHS ester (Thermo Fisher) (see the section of Preparation of fluorescently labeled cell). The symbiotic organs of the nymphs were dissected in 10 mM PBS buffer within 2-4 hours after feeding. The samples were gently placed on a coverslip, and the sample chamber was assembled with two coverslips and two pieces of double-sided tape to act as a thin channel for observation under an optical microscopy.

#### Competitive infection assay

A bacterial solution containing each 5,000 cells/μl of [*C. insecticola* Δ*flgE* + *flgE*_Ci_] (=Native flagella symbiont) and [*C. insecticola* Δ*flgE* + *flgE*_Ba_] (=Swapped flagella symbiont) were mixed, and 1 μl of the bacterial mixture was fed to the freshly molted second instar nymphs of the bean bug. The gut symbiotic organ (crypt-bearing M4 region) was dissected 48, 72, 96, and 120 h after inoculation, homogenized by a pestle, and total DNA was extracted by QIAamp DNA Mini Kit (Qiagen) and subjected to qPCR to determine bacterial titers. Due to the lack of effective antibiotic resistance markers, we decided to estimate bacterial titers in the symbiotic organ by qPCR specifically targeting *flgE*_Ci_ and *flgE_Ba_*. qPCR was conducted as previously reported^38^ with specific primer sets for *flgE_Ci_* (F[5’-TATCAGCTGTCGAACAACGG-3’] and R[5’-ATGTTGCCGTTGTCATCGAG-3’]) and *flgE_Ba_* (F[5’-CAGATTTCTCGAAACGGAGACC-3’] and R[5’-ATCGAGTTCGCGTACATGTC-3’]). By using the bacterial titers, competitive index (CI) values were calculated by (output Swapped flagella symbiont/input Swapped flagella symbiont)/(output Native flagella symbiont/input Native flagella symbiont) and statistically evaluated by the 1-sample *t* test (against CI = 1.0).

#### Statistical analysis

All statistical analyses were performed by GraphPad Prism 9.1.0 (GraphPad Software) and R ver 4.1.3^39^.

## Supplementary Text

### Numerical simulations of a bacterium swimming in a narrow tube

#### A. Problem setting and governing equations

Consider a bacterium swimming in a narrow tube of radius *R*. Assume that the tube is filled with an incompressible Newtonian fluid of density *ρ* and viscosity *μ* and that the tube length *L* is sufficiently long compared to the tube radius (*L* >> *R*). Due to the small size of bacteria, the typical particle Reynolds number can be less than 1^40^ and the inertia of the flow can be neglected. Fluid motion is then governed by the viscous dominated Stokes equation, and the flow field at an any point ***x*** is given by the following boundary integral equation^41^:

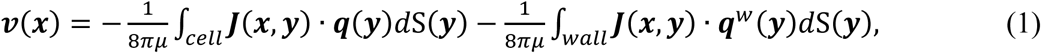

where ***q*** and ***q****^w^* are the viscous traction on the cell surface and tube wall, respectively. The first term on the right represents the flow produced by the bacterium and the second term represents the boundary conditions of the tube. ***J*** is the Green’s function of Stokeslet, which is given by

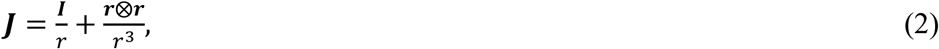

where ***r*** = ***x*** – ***y***, *r* = |***r***|, and ***I*** is the identity matrix. The bacterium is thought to swim either in the unwrapped mode, in which the flagellum helixes behind the bacterium, or in the wrapped mode, in which the flagellum wraps around the bacterium (Fig. S6b). From the state of zero inertia, it is assumed that the bacterium swims with no force and torque:

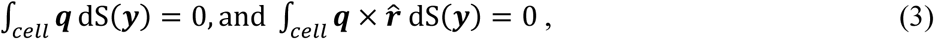

where ***r̂*** = ***y*** − ***y***_*g*_ is the relative position vector of the cell surface from the center of gravity ***y****_g_*. It is also assumed that the flagellum is driven by the motor torque ***M***_m_ at the base of the flagellum and that the torque ***M***_m_ and the fluid resistance are balanced for bacterial swimming:

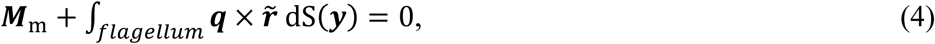

where *r̂* = ***y*** − ***y***_*g*_ is the relative position from the base of the flagellum ***y****_b_*.

#### B. Swimmer model

The bacterial body is modelled as a cylinder of length *H* with spherical ends of diameter *D* (see Fig. S6a). The ratio of *H* to *D* is set to *H*/*D* = 3 in the flow simulation. Assuming that the bacterium is moving rigidly, consider the translational swimming velocity ***U***, the rotational angular velocity of the cell body **Ω** and the angular velocity of the flagellum ***ω***:

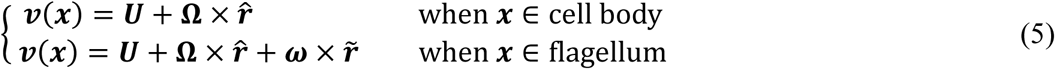

The flagellar radius *a* is sufficiently small compared to the length, and the helical shape of the flagellum is denoted by the orthonormal body frame of the center line. The flagellar surface is then modelled as a cylindrical body with radius *a* from the center line.

In the case of unwrapped mode, the flagellar center line is given by ref^42^:

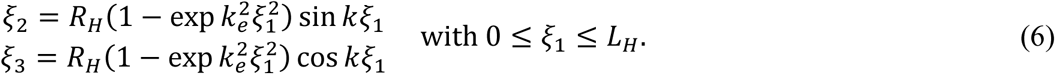

ξ_*i*_ is the orthonormal body frame with the base of the flagellum as the origin, and *R_H_* and *L_H_* are the helix radius and length, respectively. *k* is the wave number of the helix, and *k_e_* controls how quickly the helix grows to its maximum amplitude with growing distance from ξ_1_ = 0.

In the case of wrapped mode, the flagellar center line is given by the following helix equation:

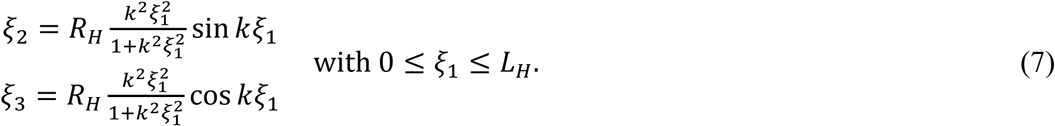

The bacterial body and flagellar surface are discretized by 5120 and 4800 triangular meshes, respectively.

#### C. Numerical procedures

To simulate the bacterial swimming, we consider resistance problems with respect to unknowns ***U***, **Ω**, ***ω***, ***q*** and ***q****^w^*. The cell surface and tube wall are discretized by triangular element and all physical quantities are computed at each vertex. Then, the boundary integral equation (1) is computed by a numerical Gaussian integration scheme^43^, and we have the following vector form of Eq. (1):

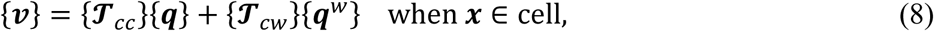

and

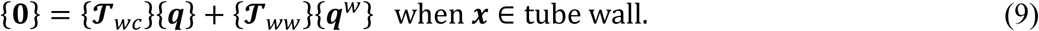

Equation (8) represents the motion of the bacterium and Eq. (9) is the no-slip boundary condition at the tube surface. The vector size of {***v***} and {***q***} are 3*N*, where *N* is the total number of nodes on the cell surface. Whereas the size of {**0**} and {***q***^*w*^} are 3*M*, where *M* is the number of nodes on the wall. Accordingly, the matrix size of {***T**cc*}, {***T***_*cw*_}, {***T***_*wc*_} and {***T***_*w*_} are 3*N* × 3*N*, 3*N* × 3*M*, 3*M* × 3*N*, and 3*M* × 3*M*, respectively. Substituting Eq. (5) into Eq. (8) and applying the force-and torque-free conditions (Eqs. 3 and 4), the system can be extended to following (3*N* + 3*M* + 9) × (3*N* + 3*M* + 9) matrix system:

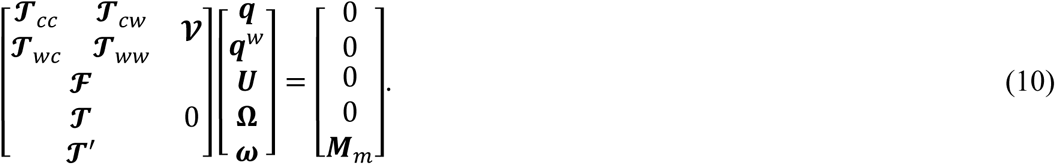

Matrix component ***F***, ***T***, and ***T***′ are computed from Eqs. (3) and (4) with the numerical Gaussian integration, and ***V*** is given by Eq. (5). The dense matrix of (10) is solved with respect to unknowns ***q***, ***q****^w^ **U***, **Ω**, and ***ω*** by a lower-upper (LU) factorization technique^43^. Once the translational velocity ***U*** and the angular velocity **Ω** and ***ω*** are given, all material points on the cell surface are updated by a second-order Runge-Kutta method. In addition, the flow field can be calculated by substituting ***q*** and ***q***^*w*^ into Eq. (1).

#### D. Numerical parameters

Flow simulations are performed in a time-space non-dimensionalised by the fluid viscosity μ, motor torque |***M****_m_*|, and body length *H*. In other words, the length, time, and viscous traction are scaled as ***x***\* = ***x***/*H*, *t*\* = *t*|***M****_m_*|/μ*H*^3^, and ***q***\* = ***q****H*^3^/|***M****_m_*|, respectively. To simulate the bacterial swimming in microchannels, the tube radius *R*/*H* is set in the range of 0.2 to 0.34 and the channel length *L* is set to *L*/*H* = 13. If we assume *H* = 2.5 μm (Fig. S6b), these values are equivalent to *R* = 0.5 to 0.85 μm and *L* = 32.5 μm.

The number of elements *Ne* is set to *Ne* = 9920 for the cell surface (*N* = 4964) and *Ne* = 10008 for the tube wall (*M* = 5040) so that typical mesh size Δ*x* is smaller than 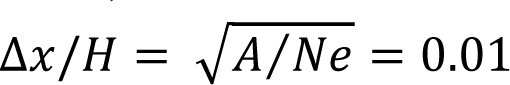, where *A* is the surface area. Accuracy of calculating the lubrication flow between the cell body and the wall depends on the mesh size. Thus, a spatial resolution of approximately 25 nm can be calculated with the mesh size of Δ*x*/*H* = 0.01.

To mimic the flagellar waveforms observed in experimental measurements, the wave number *k* is set to *kH* = 2.5π for the unwrapped mode and *kH* = 5π for the wrapped mode, respectively, and the helix radius and length are set to *R_H_*/*H* = 0.1 and *L_H_*/*H* = 2 for the unwrapped mode and *R_H_*/*H* = 0.163 and *L_H_*/*H* = 0.9 for the wrapped mode. The helix growing factor *k_e_* is also set to *k_e_H* = 1, and the flagellar radius *a* is assumed to be *a*/*H* = 0.0125^42^.

Time increment Δ*t* for the calculation is set to Δ*t*|***M***_*m*_|⁄μ*H*^3^ = 0.01. This corresponds to Δ*t* = 78 μsec by assuming μ = 1 mPa.s, and |***M***_*m*_| = 2000 pN.nm (see Supplementary text).

#### E. Flow field

The unwrapped mode flow field in free space environment produces an extensional flow in the front to back direction and a suction flow in the lateral direction (Fig. S6c). On the other hand, in the narrow tube, no lateral suction flow is generated and the fluid friction between the cell body and the wall provides strong resistance (Fig. 3g *Top*). As a result, the flow generated by the flagellum contributes only to the agitation of the fluid around the flagellum and less to its propulsion.

The wrapped mode creates a vortex flow in free space (Fig. S6c), while in the narrow channel the flagellum scrapes the fluid in the gap like a corkscrew, creating a laminar flow structure in the tube and contributing to cell propulsion (Fig. 3g *Bottom*).

#### F. Swimming velocity and angular velocity (Fig. 3h and Fig. S6d)

The swimming speed in free space without the tube is *U*_0_ μ*H*^2^⁄|***M***_*m*_| = 0.094 for the unwrapped mode. This corresponds to *U*_0_ = 30 μm/sec, whereas it is 16 μm/sec for the wrapped mode.

In the wrapped mode, the swimming speed is relatively maintained even in narrow channels. In the unwrapped mode, however, the speed decreases monotonically as smaller gap and when Δ*R/*(*H*/2) is less than 0.1, the speed is less than 4% of *U*_0_. We again assume *H* = 2.5 μm, the gap in a circular tube of 1 micro diameter corresponds to Δ*R*/(*H*/2) = 0.025 and the swimming speed is estimated to be *U* = 180 μm/min for the wrapped mode. This speed is comparable to the experimental measurement (e.g. 104 μm/min using *C. insecticola* in Table S1). In the unwrapped mode, no solution was obtained for conditions below Δ*R*/(*H*/2) = 0.1 due to computational instability.

We also calculate the rotational angular velocity in the narrow tube. Effect of small gap on the rotation is relatively small in both modes. Which indicates that the flagellar motor produces sufficient large torque to generate cellular rotation even in the narrow channel.

## Numerical simulations of flagellar dynamics and a hook stiffness

### 1. Overview of numerical simulation model

A polar flagellated bacterium studied in this work typically has a few flagella at one pole that form a bundle during propulsion. In our mechanical modeling of a flagellar wrapping, we assume the bundle of flagella as a single elastic helical filament. For the deformations of a flexible filament, we compute all the elastic modes of stretching, bending, and twisting, taking full account of the geometric nonlinearities inherent to such slender structures (Fig, S8a), based on the method reported previously^21^. The flagellar filament is connected to a rigid cell body with a flexible hook, a tiny functional mechanical element. The motor torque is transmitted to the filament without loss even when the orientation of the flagellar filament at the base deviates from the rotational axis of the motor. Because the hook is short enough (less than 100 nm) compared to a length of flagellar filament of typically several μm, we take into account effects of the hook flexibility by modelling it as a torsional spring at the connecting point with some stiffness that can be changed independently from that of the flagellar filament. As a crucial hypothesis, we assume short-range attractive interactions between distant segments of a flagellar filament. Such a weakly sticky flagellum assumption is consistent with our observation that a flagellar bundle remains stable in absence of any hydrodynamic effects at stalled rotary motor condition. Finally, the hydrodynamic force from the surrounding fluid is considered at the level of the resistive force theory, meaning that the viscous force is proportional to the local velocity with the local shape-dependent friction coefficients implying the non-local nature of the background viscous flow. In the present composite elastic-rigid model, the bacterial flagellar motor is modelled as a torque dipole embedded at the connecting point.

In the numerical simulation, a helical flagellar filament is discretized into a chain of nodes connected by short segments of approximately constant length a, while the cell body is modelled as a cylindrical rigid body of length 2*H* with spherical end caps of diameter *D* The total energy of this system, *E*_total_, consists of a number of different contributions:

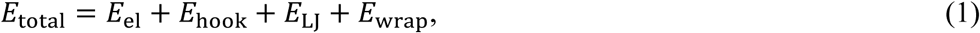

where *E*_el_ is the elastic deformation energy of the helical filament that is the sum of the stretching, bending, and twisting energies, *E*_hook_is the torsional spring potential that accounts the bending stiffness of the hook segment, *E*_LJ_ is a truncated Lennard-Jones potential that represents an attractive interaction between the distant nodes as well as the self-avoiding interaction in the filament, and *E*_wrap_ is the excluded volume interaction potential between the filament and the cell body that defines the wrapping configuration.

The forces and torques acting on each node of the flagellar filament, as well as those on the rigid cell body, can be computed via the variational methods based on the potential energy given in Eq. (1). The bacterial motion is directly driven by the internal motor torque, ***M***_*m*_ = *M*_*m*_***t***, where ***t*** defines the axis of the cell body. The flagellar filament is attached at the connecting point *x* = ***R***_*B*_ + *H**t***, where ***R***_*B*_ represents the center of mass of the cell body. In our discrete model, the motor torque on the filament is decomposed into the orbital and spinning components about the local tangent of the filament at its base, *u*. The resulting force ***f***_*m*_ and torque *T*_*m*_ are applied at the 1st node of the discrete filament. The reaction force and torque on the cell body are given by ***f***_*m*_ and −(***M***_*m*_ + *Ht* × ***f***_*m*_). We have confirmed both analytically and numerically that all the internal forces and torques, including the motor produced forces and torques, satisfy the action-reaction law, ensuring the overall force and torque free conditions for a freely swimming bacterium. At low Reynolds number relevant to our problem, the positions and orientations of each node of the filament evolve according to the overdamped Langevin equations that describe the translational and rotational motions of the local director frame attached at each node in presence of thermal forcing. Similarly, the viscous force and torque balance equations for the cell body describe the translation of its center of mass position ***R***_*B*_and the rotation of its body-fixed frame.

We use the Euler iteration method to numerically integrate the appropriately rescaled equations of motion with a nondimensional time step of typically 0.01. The output values are calculated every 10^5^−10^6^ steps, and the total simulation time is 10^7^−10^8^ steps. All the nondimensional parameters in the equations are matched with those directly relevant to our experimental observations. Values of some key parameters that may change qualitive behaviors of our numerical model are further explained below. A more detailed description of our mathematical biomechanical modeling will be reported elsewhere.

### 2. Relevant parameters

A flagellar filament is assumed to have a uniform radius *R* and pitch *P* in its natural configuration. In the swimming mode, we set *R_normal_* = 0.2 μm, *P_normal_* = 2.0 μm, valid to the Normal form, with the three full turns along the arclength. In the wrapped mode, we set *R_coil_* = 0.6 μm, *P_coil_* = 1.4 μm, valid to the Coil form, with the two full turns along the arclength. Note that the both forms are left-handed helices.

We denote the bending, twisting, and stretching moduli of a flagellar filament as *A*, *C* and *K*, respecively. The bending modulus of a bacterial flagellum *A* has been measured in the previous studies, which on the whole suggest *A* = 2.3-3.5 pN · μm^220^. In this study, we assume *A* = 3.0 pN · μm^2^ per a filament. In contrast, the twisting modulus *C* and the stretching modulus *K* have not been measured directly. Assuming an isotropic elastic rod of diameter *a*, we choose *Ka*^2^/*A* = 16, which ensures that segment length variations are negligibly small so that the filament is regarded approximately inextensible. For an isotropic elastic rod, we have *C*/*A* = 1/(1 + *v*), where *v* is the Poisson’s ratio of the rod material. Since 0 ≤ *v* ≤ 1/2 for ordinary materials, we usually assume 0 ≤ *C*/*A* ≤ 1. Nevertheless, considering the complexities of the supramolecular structures of a flagellar filament, *C*/*A* is allowed to take a wider range of values. We therefore change the twist/bend ratio *C*/*A* between 0.5 to 1.0.

On the other hand, we have determined the stiffness of the hook bending torque as 3.36 pN · μm for *C*. *insecticola* and 16.7 pN · μm for *B*. *anthina*. Assuming that the bundle analyzed consisted of three identical filaments, we have the stiffness per hook as *k*_θ_ = 1.12 pN · μm ·μm for *C*. *insecticola* and *k*_θ_= 5.54 pN · μm for *B*. *anthina*. The hook bending modulus, *A*_hook_, is related to the stiffness *k*_θ_ determined from the main axis angle Brownian motion, as *A*_hook_= *k*_θ_*l*, where *l* is the hook length. Assuming *l* ≈ 60 *nm*, we find the rescaled hook stiffness as *A*_hook_/*A* = 0.02 for *C*. *insecticola* and 0.11 for *B*. *anthina*. In the simulations, we change *A*_hook_/*A* from 0.02 to 0.15.

We set the magnitude of torque generated by the flagellar rotary motor as ***M***_*m*_ = 0.40-0.50*A*/*R_coil_*. Assuming *A* = 3.0 pN · μm^2^ and *R_coil_*= 0.6 μm, it amounts to ***M***_*m*_ = 2000-2500 pN · nm per rotary motor, which is only slighly higher than the previously reported values 1300-1800 pN · nm for *E. coli*^20^.

### 3. Results

In Fig. S8b-d, we show the three qualitatively distinctive wrapping behaviors observed in the simulations for varying hook/filament stiffness ratio, *A*_hook_/*A*, and the twist/bend stiffness ratio of filament, *C*/*A* (Table S2).

For *A*_hook_/*A* = 0.02 and *C*/*A* = 0.75, we find a smooth wrapping transition (Movie S13), in good agreement with the experimental observation (Fig. 4a). For the increased hook stiffness *A*_hook_/*A* = 0.14 and *C*/*A* = 0.75, the wrapping behavior is distinctively different from the soft hook case, where the filament overlaps near the end of the cell body (Movie S14). The tendency is overall unchanged for *C*/*A* = 1.0. Physically, when the hook is sufficiently stiff, the filament folds before completely wrapping around the cell body, leading to the peculiar morphology dictated probably by the balance between the elasticity and intra-filament attractions. For even smaller twist/bend stiffness ratio given by *C*/*A* = 0.5 (with the stiff hook of *A*_hook_/*A* = 0.14), this feature is more evident, where the filament forms a ring without wrapping around the cell body at all (Movie S15).

These numerical results are all consistent with the optical microscopy observations, which validates our physical modeling of the wrapping bacterium. As a crucial check, the assumption of the sticky flagellar filament will have to be examined in future studies. Overall, our biomechanical modeling and its numerical simulations suggest a potentially important role of the hook stiffness upon the wrapping ability of polar flagellated bacteria.

## Supplementary Figures

**Fig. S1.**
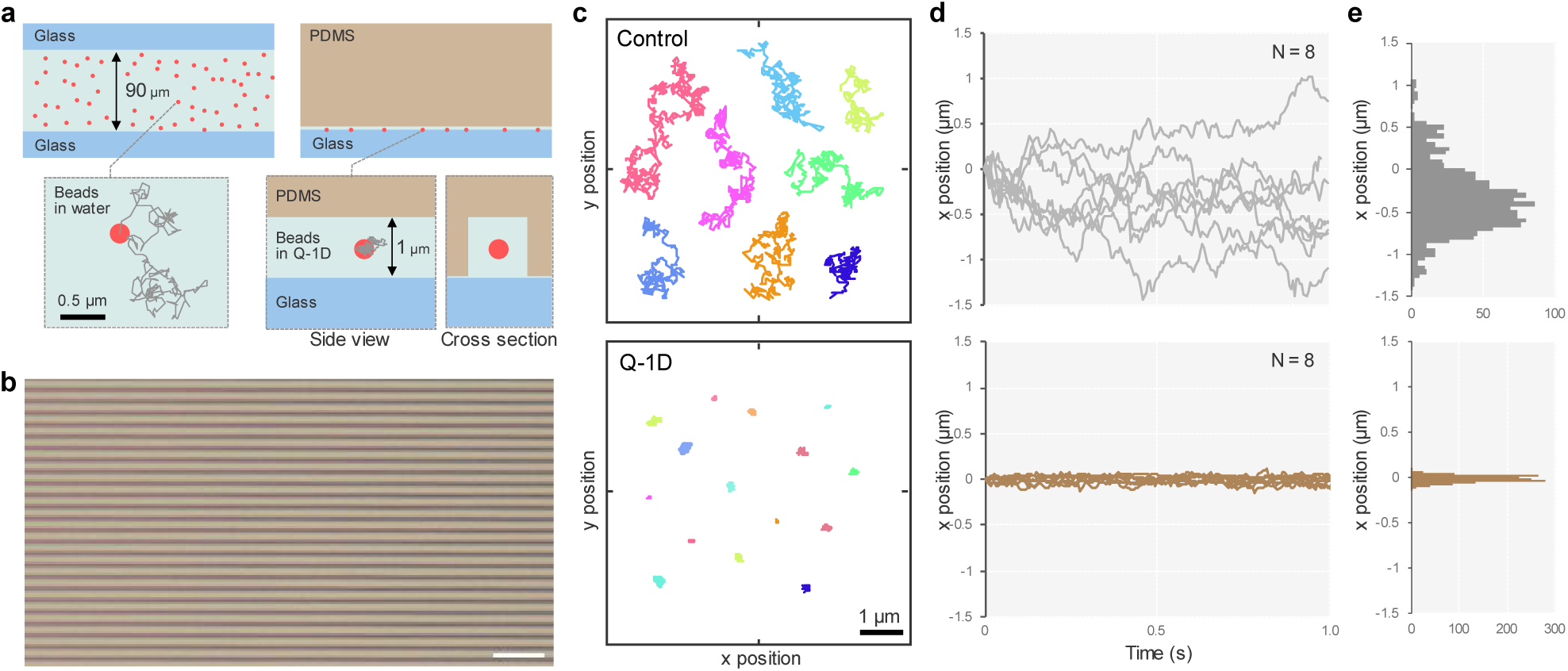
Brownian motion of microbeads in Q-1D. **a**, Confinement of microbeads. Schematic of the sample observation from a side view. Fluorescent beads with a diameter of 200 nm suspended in water were used as the sample. *Left*: Standard chamber. The chamber was assembled with two pieces of coverslip at a height of 90 µm. *Right*: Q-1D. The sample was confined in the narrow 1×1 µm square tube passage by pressing the device made by PDMS from the top. **b**, Phase-contrast image of Q-1D. Line-patterned white area is a space for the sample confinement. Scale Bar, 10 µm. **c**, Trajectories of microbeads at 5.6 ms-intervals for 1 s. **d**, Time course of the microbeads for 1 sec. *Top*: Control as standard chamber. *Bottom*: Q-1D. **e**, Distribution of the bead displacement. *Top*: standard chamber. *Bottom*: Q-1D. The variance of the bead displacement in standard chamber and Q-1D are measured as 0.313 and 0.041, respectively.

**Fig. S2.**
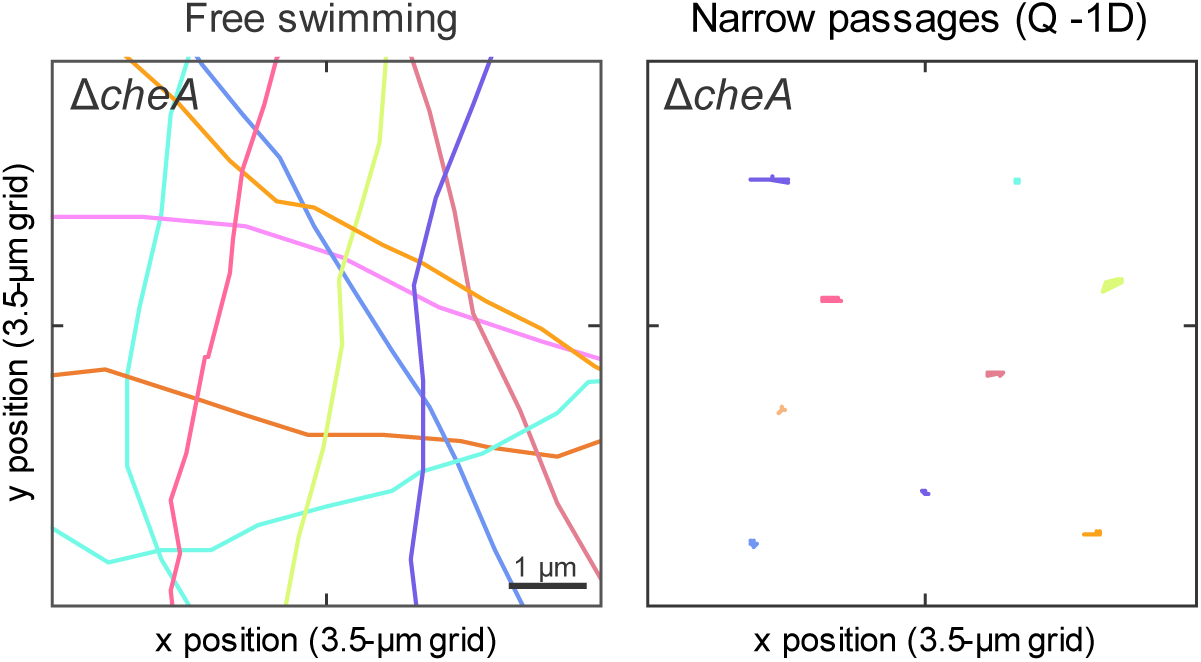
Cell behavior of Δ*cheA* deletion mutant of *C. insecticola* in Q-1D. Trajectories of the cell at 50 ms-intervals for 1 s. (*Left*) Free swimming in the standard chamber. The chamber was assembled with two pieces of coverslip at a height of 90 µm. (*Right*) Cell behavior in the narrow passages of Q-1D.

**Fig. S3.**
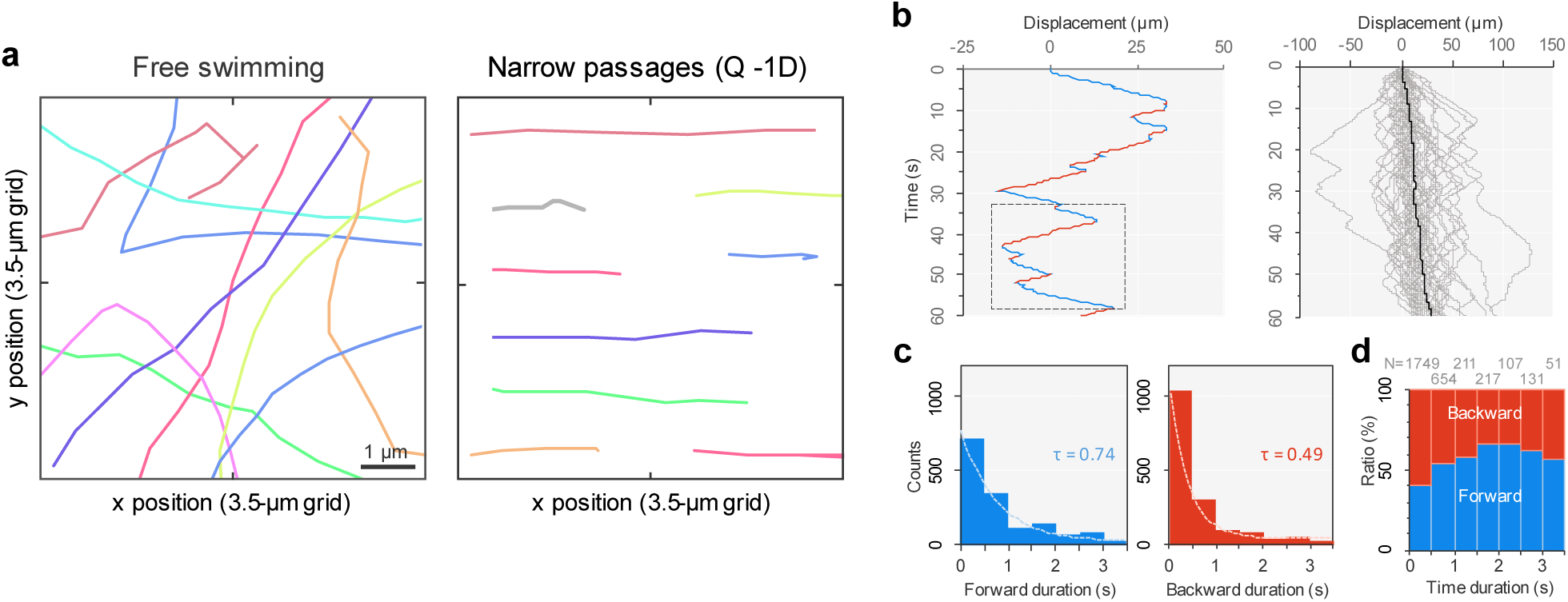
Cell behavior of *S. enterica* in Q-1D. **a**, Trajectories of the cell for 1 s in the standard chamber (*Left*) and narrow passages of Q-1D (*Right*). **b**, Time course of cell displacement in Q-1D. *Left*: Single cell. Forward and backward movements are represented as blue and red colored lines respectively. *Right*: Overlays of 50 cells and the average. **c**, Distribution of the forward and backward time duration. The duration from a directional change to the next were measured in Q-1D. Color code the same way as **b**. Dashed lines show the fit of single exponential decay, where time constant τ is presented. **d**, Ratio of the time duration of forward and backward direction. The ratio was sorted by 0.5 s, and each ratio at the time points were presented.

**Fig. S4.**
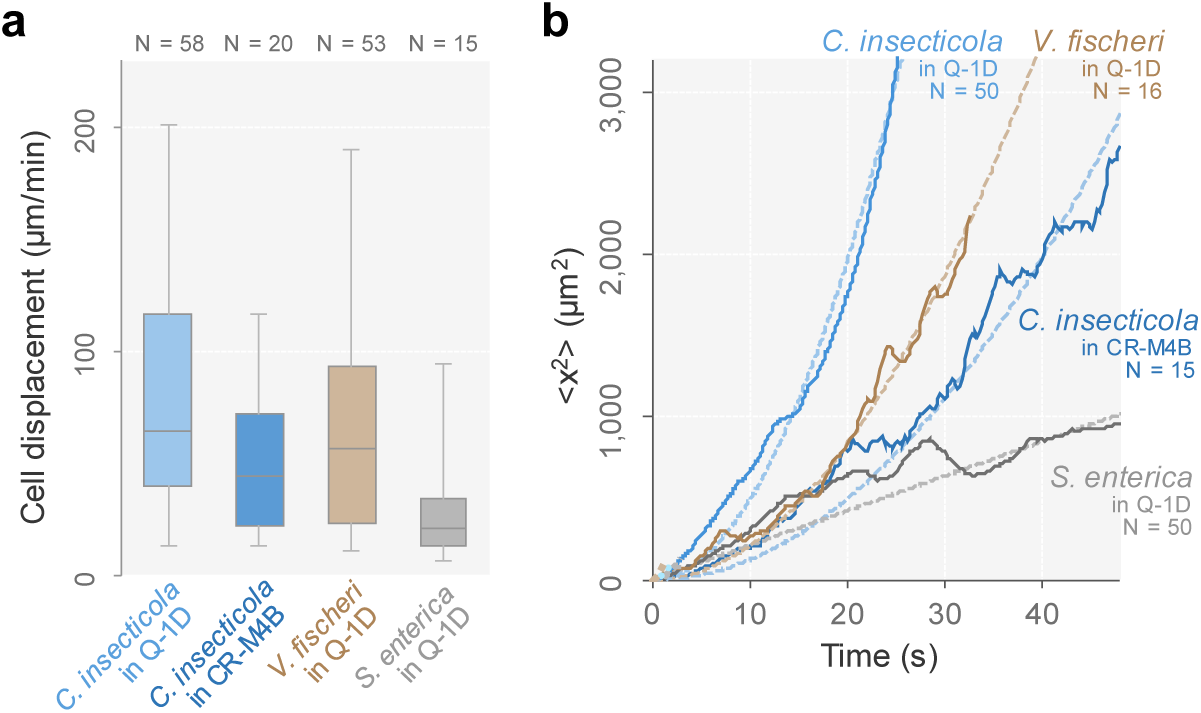
Cell displacement in a narrow passage. **a**, Net displacement of *C*. *insecticola* in Q-1D, *C*. *insecticola* in CR-M4B region in host (the same data set from Fig. 1d), *V. fischeri* in Q-1D, and *S*. *enterica* in Q-1D for 1 min. Box plot presents the minimum, maximum, sample median, and the first and third quartiles. **b**, MSD plots of cell movement. Dashed lines of *C. insecticola* and *V. fischeri* represents a hyperbolic fitting, <*x*^2^> = a*t*^2^. Dashed lines of *S. enterica* represents a linear fitting, <*x*^2^> = a*t*.

**Fig. S5.**
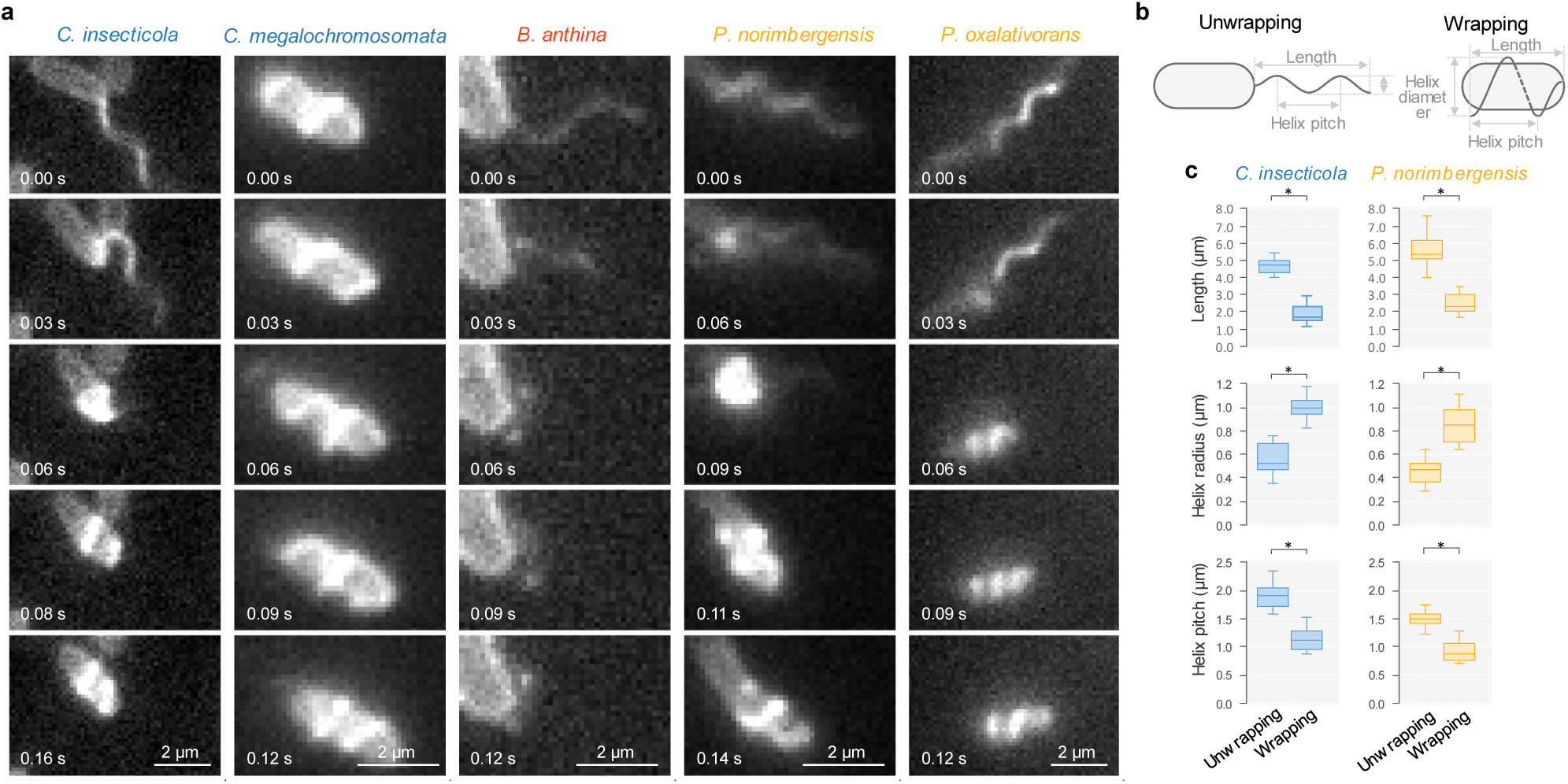
Visualization of flagellar filament in swimming cells of *Burkholderia sensu lato* group. **a**, Sequential fluorescent images of *C*. *insecticola*, *C*. *megalochromosomata*, *B*. *anthina*, *P*. *norimbergensis*, and *P*. *oxalativorans*. The swimming mode was changing from unwrapping to wrapping in *C*. *insecticola*, *P*. *norimbergensis*, and *P*. *oxalativorans*. The *C*. *megalochromosomata* cell propels itself with flagellar wrapping. The *B. anthina* cell is captured at incomplete flagellar wrapping when the flagellar filaments are folded as a ring at the proximal end. All observations are performed in the presence of 0.5% MC. **b**, Schematic of the parameter of the helix morphology in unwrapping (*right*) and wrapping cells (*left*). **c**, Measurements of the helix morphology of flagellar filaments in *C*. *insecticola*, *P*. *norimbergensis*. Box plots of flagellar length along the helix axis, a helix radius, and a helix pitch. These parameters were measured based on the fluorescent image of their flagellar filament in Q-1D (N = 20 cells). Statistical analysis was performed using the unpaired *t* test (*P* < 0.05).

**Fig. S6.**
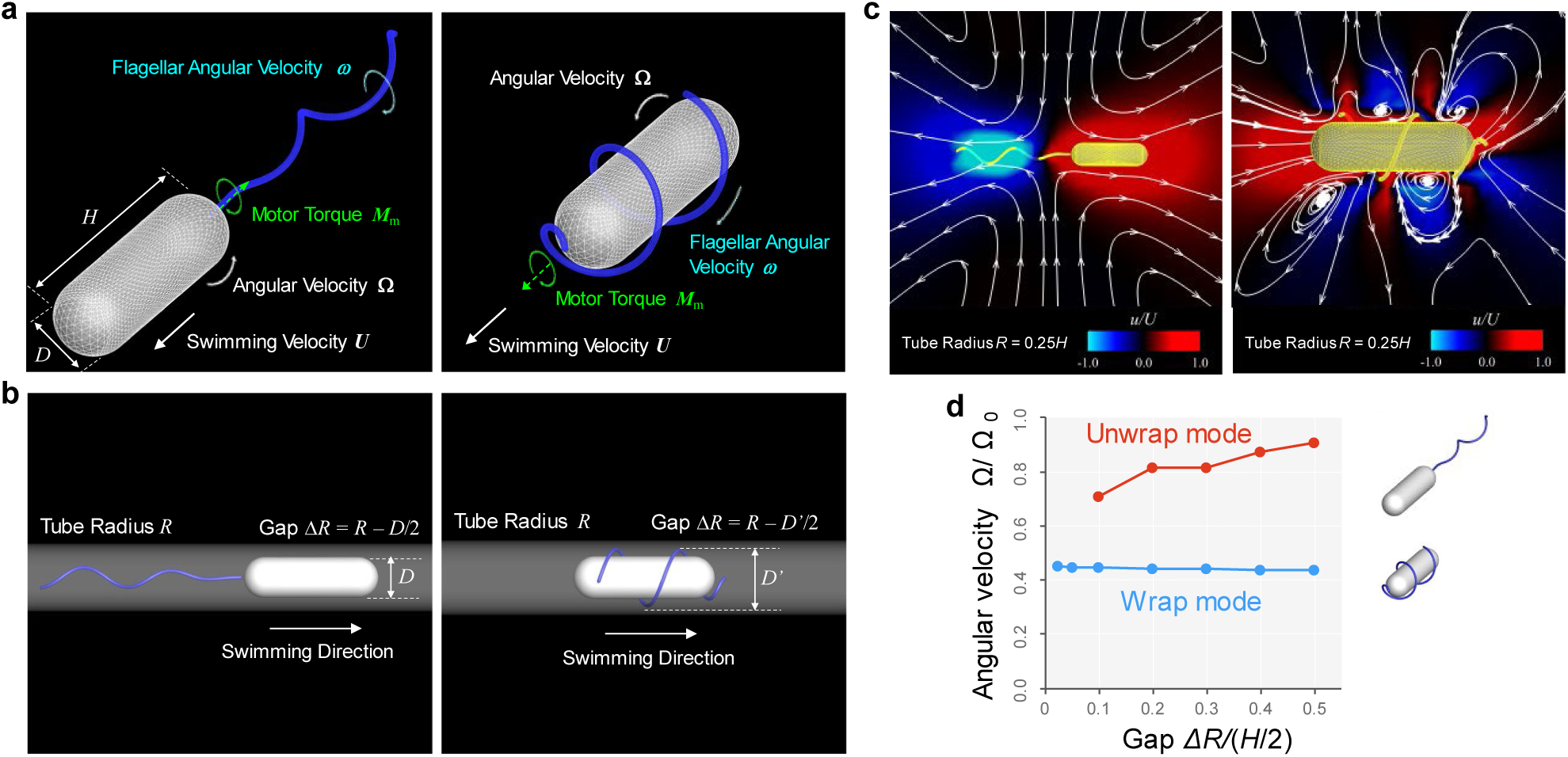
Numerical simulations of a bacterium swimming in a narrow tube. **a**, Fluid mechanical model of swimming bacteria exhibiting two distinct modes. *Left*: Unwrapped mode. *Right*: Wrapped mode. The cell body is modelled as a cylinder of length *H* with spherical ends of diameter *D*. The flagellum has a helical shape with the helical diameter and pitch set to match experimental measurements (Fig. S5bc). The flagellum is assumed to be driven by the motor torque ***M***_m_ at the root of the flagellum, and the rigid body translation velocity ***U*** and the angular velocities **Ω**, and ***ω*** are derived from the boundary integral equation of the Stokes flow. The cell body and the flagellar surface are discretized with 5120 and 4800 triangular meshes, respectively. **b**, Problem setting of the numerical simulation. *Left*: Unwrapped mode. *Right*: Wrapped mode. The bacterial model in a circular tube of radius *R* swims in one direction and the minimum distance between the wall and the bacterium is defined as Δ*R*. **c**, Time-averaged flow field around the bacterial model in free space. *Left*: Unwrapped mode. *Right*: Wrapped mode. White arrows are the streamline and contour color indicate the velocity component in the swimming direction normalized by the swimming velocity *U*. **d**, Angular velocity Ω as a function of Δ*R*. Ω_0_ is the angular velocity of the unwrap mode in a free space.

**Fig. S7.**
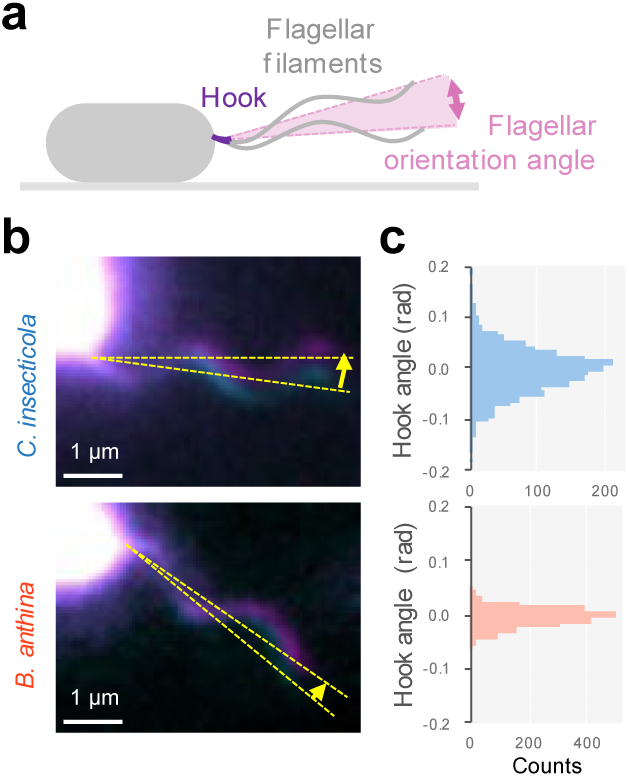
Fluctuation of flagellar filaments at the proximal part. **a**, Schematic. Flagellar rotation was inactivated by CCCP, and the cell was immobilized on a glass surface. **b**, Fluorescent images. Two sequential images are colored by magenta and cyan, respectively, and merged to see the fluctuation. Dashed yellow line is the helix axis of the flagellar filaments. **c**, Distribution of the flagellar orientation angle for a single cell recorded at 5 ms for 10 s. Hook flexibility is measured by the variance of the flagellar orientation angle (see Supplementary text). Homogeneity of variances was analyzed by Bartlett test (*P* < 0.05).

**Fig. S8.**
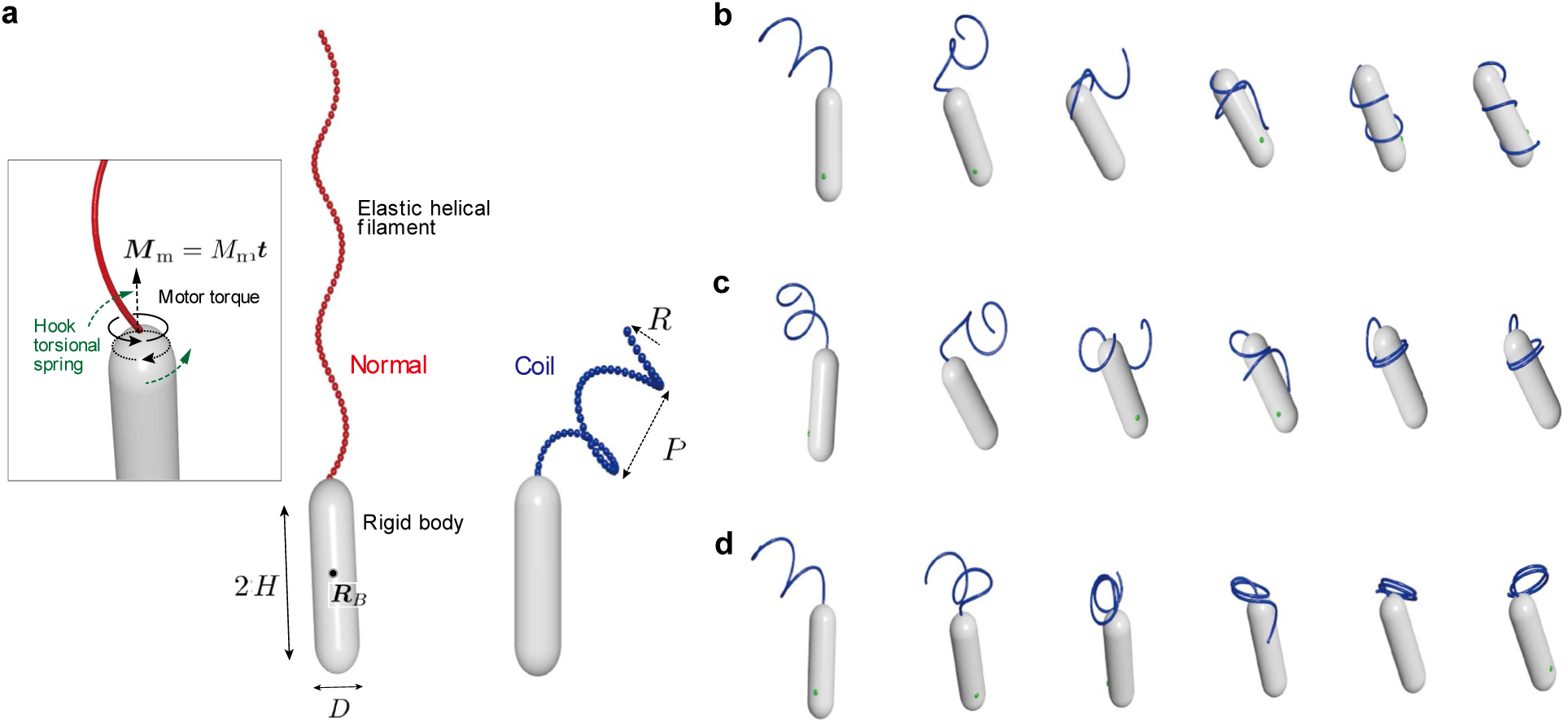
Numerical simulations of flagellar dynamics and a hook stiffness. **a**, Numerical simulation model. Schematics of the model for wrapping bacterium and some relevant geometric parameters. A set of discrete points representing a flagellar filament is explicitly drawn here. The number of the nodes is set *N* = 60. Inset shows a close-up view of the connecting point, where the torques by the flagellar motor and by the hook stiffness apply to the filament base. Note that reaction torques also apply to the cell body. **b-d**, Typical snapshots from our numerical simulations. For all three cases, the attractive interaction between distant segments in the filament is assumed, with the Lennard-Jones potential parameters ɛ_LJ_ = 20*k*_*B*_*T* and *r*_cutoff_ = 2*a*, where *k*_*B*_*T* is the thermal energy. **b**, Normal wrapping for *C*/*A* = 0.75 and *A*_hook_/*A* = 0.02 (soft hook). **c**, Incomplete wrapping for *C*/*A* = 0.75 and *A*_hook_/*A* = 0.14 (stiff hook). **d**, Ring formation for *C*/*A* = 0.50 and *A*_hook_/*A* = 0.14 (stiff hook). Note that the cell body also translates during the wrapping, but its positions are shifted to align for the visualization purpose here.

## Supplementary Tables

**Table S1.** Cell motility of 13 species in *Burkholderia* sensu lato and allied groups

| Species | Swimming speed in standard chamber ($\mu\text{m/s}$)<sup>†</sup> | Cell displacement in Q-1D ($\mu\text{m/min}$)<sup>‡</sup> | Infection rate (%)<sup>§</sup> | Flagellar wrapping<sup>¶</sup> |
| --- | --- | --- | --- | --- |
| *Caballeronia insecticola* | 26.4 ± 8.4 | 104 ± 52 | 100 | Yes |
| *Caballeronia megalochromosomata* | 22.4 ± 5.8 | 132 ± 92 | 100 | Yes |
| *Caballeronia glathei* | 22.1 ± 9.5 | 86 ± 47 | 100 | n.d. |
| *Caballeronia telluris* | 19.6 ± 6.9 | 42 ± 29 | 100 | n.d. |
| *Paraburkholderia tropica* | 34.1 ± 12.4 | 114 ± 83 | 100 | n.d. |
| *Paraburkholderia mimosarum* | 28.0 ± 6.0 | 0 ± 0 | 0 | n.d. |
| *Paraburkholderia tuberum* | 23.0 ± 5.3 | 0 ± 0 | 0 | n.d. |
| *Paraburkholderia graminis* | 34.1 ± 10.5 | 78 ± 58 | 100 | n.d. |
| *Burkholderia plantarii* | 23.4 ± 7.9 | 6 ± 6 | 0 | n.d. |
| *Burkholderia anthina* | 14.7 ± 4.8 | 0 ± 0 | 10 | No |
| *Pandoraea oxalativorans* | 20.9 ± 7.6 | 43 ± 27 | 100 | Yes |
| *Pandoraea norimbergensis* | 19.8 ± 3.8 | 61 ± 35 | 100 | Yes |
| *Cupriavidus taiwanensis* | 21.7 ± 9.3 | 0 ± 0 | 0 | n.d. |
| *Salmonella enterica* | 22.8 ± 5.2 | 25 ± 10 | n.d. | No |

<sup>†</sup>Swimming speed was measured in the growth medium of each bacterium.
<sup>‡</sup>Cell displacement was measured in the growth medium with 0.4% MC.
<sup>§</sup>Infection rate was measured by the number of infected stink bugs/total number of investigated stink bugs<sup>7</sup>.
<sup>¶</sup>Flagellar wrapping was examined with the species that can be labeled the flagellar filaments by amine-reactive fluorescent dye. The species that cannot visualized the flagellar filaments were presented as not detected (n.d.).

**Table S2.** Details of the simulation data

| Mode | $A_{\text{hook}}/A^{\dagger}$ | $C/A^{\ddagger}$ | Motor | Flagellar form | Movie |
| --- | --- | --- | --- | --- | --- |
| Wrapping | 0.02 (soft) | 0.75 | 0.40 CW | Coil | Movie S13 |
| Incomplete wrapping | 0.14 (stiff) | 0.75 | 0.40 CW | Coil | Movie S14 |
| Ring formation | 0.14 (stiff) | 0.50 | 0.40 CW | Coil | Movie S15 |

$^{\dagger} A_{\text{hook}}/A$ : the ratio of the hook bending stiffness to that of the flagellum. $^{\ddagger} C/A$ : twist to bend ratio of flagellum. Motor torque $\mathbf{M}_m$ is shown in a unit of $A/R_{\text{coil}}$ . All the other simulation parameters will be provided in Supplemental Text. Attraction parameter (LJ potential): $\epsilon_{\text{LJ}} = 20k_B T$ , $r_{\text{cutoff}} = 2a$ .

**Table S3.** Culture condition of bacterial species in this study

| Species | Medium | Temperature ( °C) |
| --- | --- | --- |
| *Caballeronia insecticola* (RPE64) JCM 31142 | YG | 28 |
| *Caballeronia megalochromosomata* DSM 100850 | YG | 28 |
| *Caballeronia glathei* JCM 10563 | YG | 28 |
| *Caballeronia telluris* LMG 22936 | YG | 28 |
| *Paraburkholderia tropica* DSM 15359 | YG | 28 |
| *Paraburkholderia mimosarum* DSM 21841 | YG | 28 |
| *Paraburkholderia tuberum* DSM 18489 | YG | 28 |
| *Paraburkholderia graminis* DSM 17151 | YG | 28 |
| *Burkholderia plantarii* JCM 5492 | YG | 28 |
| *Burkholderia anthina* DSM 16086 | YG | 28 |
| *Pandoraea oxalativorans* DSM 23570 | YG | 28 |
| *Pandoraea norimbergensis* JCM 10565 | YG | 28 |
| *Cupriavidus taiwanensis* DSM17343 | YG | 28 |
| *Salmonella enterica* serovar Typhimurium strain SJW1103 | LB | 37 |
| *Vibrio fischeri* (*Aliivibrio fischeri*) ATCC7744 | SWT | 28 |

**Table S4.**
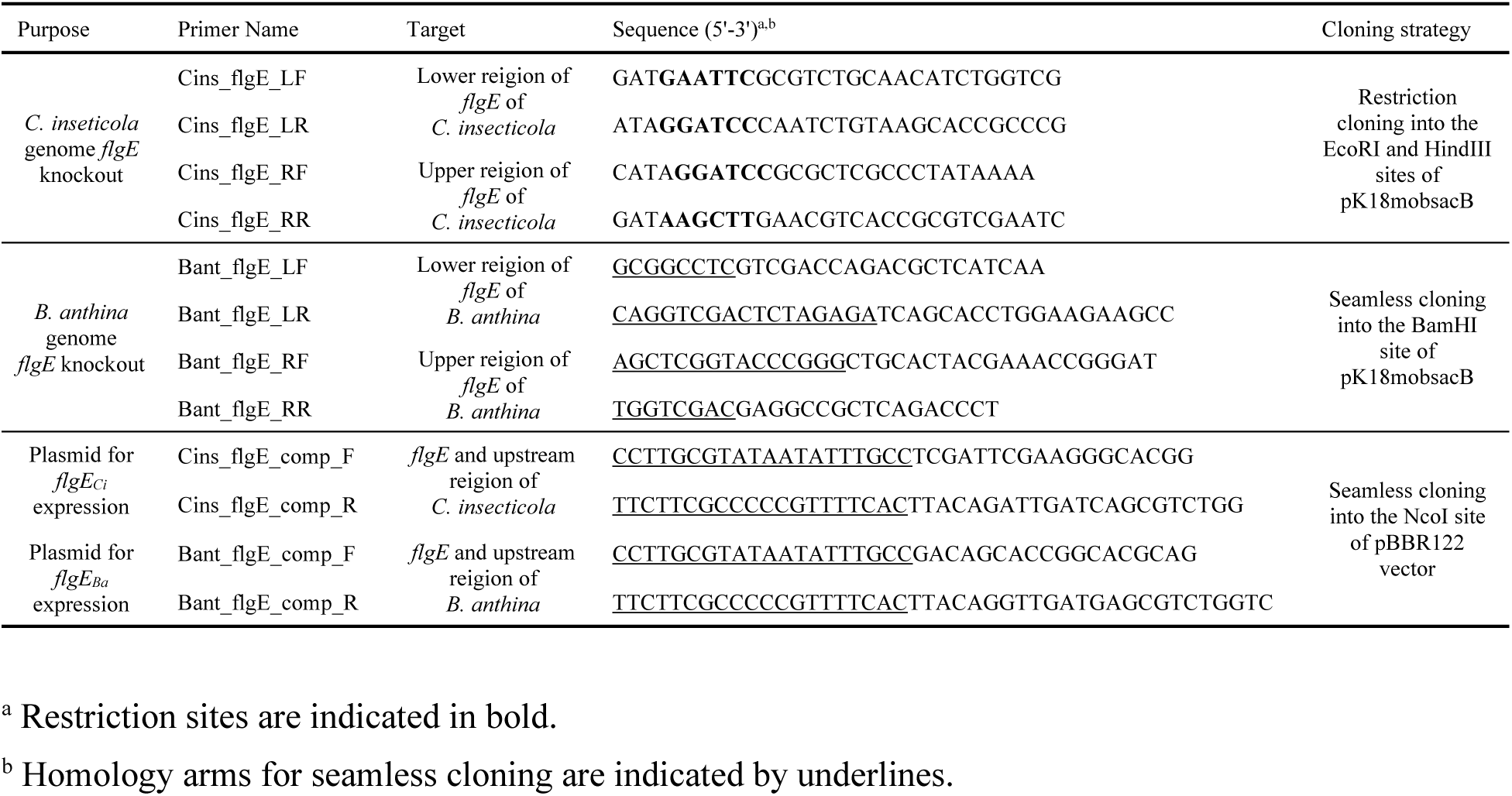
Culture condition of bacterial species in this study

**Table S5.** Strains and mutants used in this study

| Strains | Description | Reference |
| --- | --- | --- |
| *Caballeronia insecticola* RPE64 | Wild type | Ref<sup>8</sup> |
| *C. insecticola* $\Delta cheA$ | Chemotaxis deletion mutant. No switchback. | Ref<sup>10</sup> |
| *C. insecticola* GFP | GFP expressing cell | Ref<sup>37</sup> |
| *C. insecticola* $\Delta flgE$ | Gene coding hook protein deletion mutant | This study |
| *C. insecticola* $\Delta flgE + flgE_{Ci}$ | Complement with *C. insecticola flgE* | This study |
| *C. insecticola* $\Delta flgE + flgE_{Ba}$ | Complement with *B. anthina flgE* | This study |
| *Burkholderia anthina* DSM 16086 | Wild type | Ref<sup>7</sup> |
| *B. anthina* $\Delta flgE$ | Gene coding hook protein deletion mutant | This study |
| *B. anthina* $\Delta flgE + flgE_{Ba}$ | Complement with *B. anthina flgE* | This study |
| *B. anthina* $\Delta flgE + flgE_{Ci}$ | Complement with *C. insecticola flgE* | This study |

## Supplementary Movies

**Movie S1**. Behavior of *C*. *insecticola* in the CR of *R. pedestris*. The second instar nymphs were fed with a suspension of GFP expressing *C*. *insecticola*. The symbiontic organs of the nymphs were dissected 2 hours after feeding. M4 region is located at the right of the movie.

**Movie S2**. Flagellar dynamics of WT *C*. *insecticola* cells in the midgut of *R. pedestris*. The second instar nymphs were fed a suspension of symbiotic bacteria cells with fluorescently labeled body and flagellar filaments. The symbiontic organs of the nymphs were dissected 2 hours after feeding. M4 region is located at the upper right of the movie. Yellow arrows indicate flagellar wrapping cells. Area 26.0 μm × 19.5 μm.

**Movie S3**. Cell behavior of wild type *C*. *insecticola* in Q-1D captured with phase-contrast microscopy. Narrow passages of the Q-1D were align parallel to the horizontal axis. Area 300 μm × 300 μm.

**Movie S4**. Cell behavior of Δ*cheA* mutant of *C*. *insecticola* in Q-1D captured with phase-contrast microscopy. Area 300 μm × 300 μm.

**Movie S5**. Cell behavior of *S*. *enterica* in Q-1D captured with phase-contrast microscopy. Area 300 μm × 300 μm.

**Movie S6**. Cell behavior of *V*. *fischeri* in Q-1D captured with phase-contrast microscopy. Area 300 μm × 300 μm.

**Movie S7**. Cell behavior of *P*. *norimbergensis* in Q-1D captured with phase-contrast microscopy. Area 300 μm × 300 μm.

**Movie S8**. Cell behavior of *B*. *anthina* in Q-1D captured with phase-contrast microscopy. Area 300 μm × 300 μm.

**Movie S9**. Cell behavior and flagellar dynamics of 5 species of *Burkholderia sensu lato* group: *C*. *insecticola*, *C*. *megalochromosomata*, *B*. *anthina*, *P*. *norimbergensis*, and *P*. *oxalativorans*. Cells were labeled by fluorescent dye, suspended in the liquid medium containing 0.5% MC and captured with fluorescent microscopy at 5-ms interval. Area 35.1 μm × 31.2 μm.

**Movie S10**. Visualization of flagellar filaments of *C*. *insecticola* cell in Q-1D. Cells were labeled by fluorescent dye, suspended in the liquid medium containing 0.4% MC and captured with fluorescent microscopy at 12.5-ms interval. Area 32.9 μm × 5.2 μm.

**Movie S11**. Comparison of the dynamics of flagellar filaments in Q-1D. The cells of *C*. *insecticola* WT and Δ*cheA* mutant cells were labeled by fluorescent dye, suspended in the liquid medium containing 0.4% MC and captured with fluorescent microscopy. Area of each real movie 32.9 μm × 5.2 μm.

**Movie S12**. Fluctuation of flagellar filaments. Flagellar rotation was inactivated by CCCP, and the cell was immobilized on a glass surface. Flagellar filaments were wobbled at the proximal end with small angle due to Brownian motion. Cells were labeled with amine-reactive fluorescent dye to see the flagellar filaments, and captured at 5-ms interval. Area 10.4 μm × 3.9 μm.

**Movie S13**. Numerical simulations of flagellar wrapping. The attractive interaction between distant segments in the filament is assumed, with the Lennard-Jones potential parameters ɛ_LJ_ = 20*k*_*B*_*T* and *r*_cutoff_ = 2*a*, where *k*_*B*_*T* is the thermal energy. The parameters of *C*/*A* = 0.75 and *A*_hook_/*A* = 0.02 are used as a soft hook.

**Movie S14**. Numerical calculation of incomplete flagellar wrapping. The attractive interaction between distant segments in the filament is assumed, with the Lennard-Jones potential parameters ɛ_LJ_ = 20*k*_*B*_*T* and *r*_cutoff_ = 2*a*, where *k*_*B*_*T* is the thermal energy. The parameters of *C*/*A* = 0.75 and *A*_hook_/*A* = 0.14 are used as a stiff hook.

**Movie S15**. Numerical calculation of flagellar ring formation. The attractive interaction between distant segments in the filament is assumed, with the Lennard-Jones potential parameters ɛ_LJ_ = 20*k*_*B*_*T* and *r*_cutoff_ = 2*a*, where *k*_*B*_*T* is the thermal energy. The parameters of *C*/*A* = 0.50 and *A*_hook_/*A* = 0.14 are used as a stiff hook.

**Movie S16**. Comparison of numerical calculation and real images. Numerical results for rigid and flexible hook (see also Movie S13 and Movie S15) were presented with the dynamics of flagellar filaments in *C. insecticola* and *B. anthina*. The cells were labeled by fluorescent dye, suspended in the liquid medium containing 0.5% MC, and flagellar filaments were captured with fluorescent microscopy at 5-ms interval for 0.5 s (see also Movie S9). Area of each real movie 7.80 μm × 5.85 μm.

**Movie S17**. Visualization of flagellar filaments of *B. anthina* with FlgE_Ci_. Cells were labeled by fluorescent dye, suspended in the liquid medium containing 0.4% MC and captured with fluorescent microscopy at 5-ms interval. Area 10.4 μm × 7.8 μm.

## Notes

### Competing Interest Statement

The authors have declared no competing interest.

